# Molecular basis for the regulation of human glycogen synthase by phosphorylation and glucose-6-phosphate

**DOI:** 10.1101/2021.11.12.468446

**Authors:** Thomas J. McCorvie, Paula M. Loria, Meihua Tu, Seungil Han, Leela Shrestha, D. Sean Froese, Igor M. Ferreira, Allison P. Berg, Wyatt W. Yue

## Abstract

Glycogen synthase (GYS1), in complex with glycogenin (GYG1), is the central enzyme of muscle glycogen biosynthesis, and its inhibition has been proposed as a therapeutic avenue for various glycogen storage diseases (GSDs). GYS1 activity is inhibited by phosphorylation of its N- and C-termini, which can be relieved by allosteric activation of glucose-6-phosphate. However, the structural basis of GYS1 regulation is unclear. Here, we present the first cryo-EM structures of phosphorylated human GYS1 complexed with a minimal interacting region of GYG1 in the inhibited, activated, and catalytically competent states at resolutions of 3.0-4.0 Å. These structures reveal how phosphorylations of specific N- and C- terminal residues are sensed by different arginine clusters that lock the GYS1 tetramer complex in an inhibited state via inter-subunit interactions. The allosteric activator, glucose-6-phopshate, promotes a conformational change by disrupting these interactions and increases flexibility of GYS1 allowing for a catalytically competent state to occur when bound to the sugar donor UDP-glucose. We also identify an inhibited-like conformation that has not transitioned into the activated state, whereby the locking interaction of phosphorylation with the arginine cluster impedes the subsequent conformational changes due to glucose-6-phosphate binding. Finally, we show that the PP1 phosphatase regulatory subunit PPP1R3C (PTG) is recruited to the GYS1:GYG1 complex through direct interaction with glycogen. Our results address long-standing questions into the mechanism of human glycogen synthase regulation.

---

Glycogen serves as the main carbohydrate store and energy reserve across animal phyla and contains up to 55,000 glucose units linked by α-1,4 and α-1,6 glucosidic bonds^1^. Glycogen biosynthesis is catalyzed by the concerted actions of three enzymes in eukaryotes: (i) glycogenin (GYG, EC 2.4.1.186), which forms a short primer through stepwise attachment of glucose units onto itself^2^; (ii) glycogen synthase (GYS, EC 2.4.1.11), which “strings” glucose units to elongate the GYG-attached primer^3^; and (iii) glycogen branching enzyme (GBE, EC 2.4.1.18), which introduces branch points to a linear chain via α-1,6 linkages^4^ (Fig. 1b). In mammals, glycogen is primarily stored in the liver (for regulating glucose homeostasis during fasting) and muscle (as an energy reserve during exercise).

**Fig. 1.**
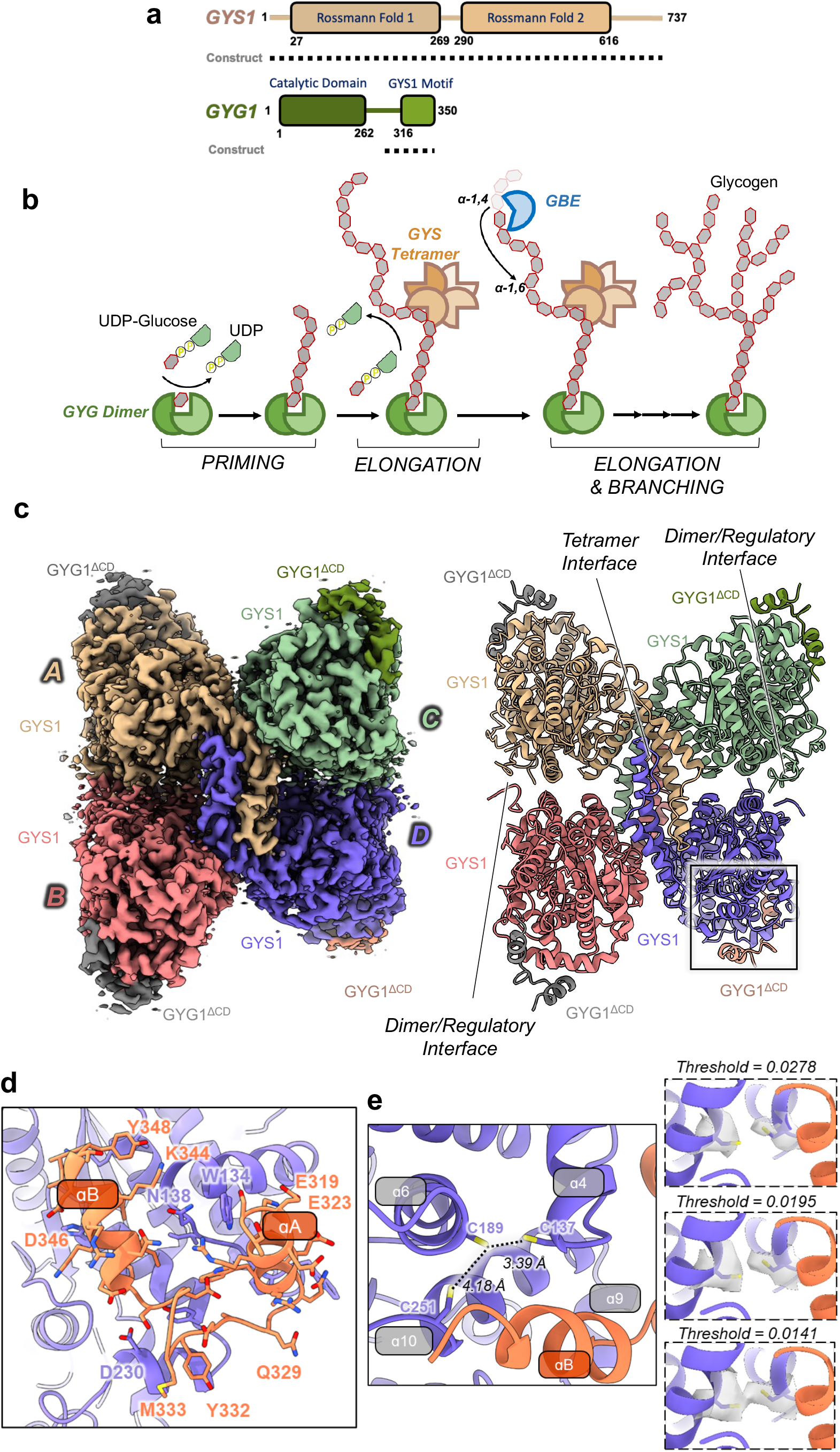
Structure of the phosphorylated inhibited/T state GYS1:GYG1^ΔCD^ complex. **a**, Domain diagrams of human GYS1 and GYG1. Dotted lines represent the construct boundaries of the GYS1:GYG1^ΔCD^ complex used in all cryo-EM experiments. **b**, Schematic of the enzymatic catalyzed reactions of GYG1, GYS1, and GBE. Glycogen synthesis is multistep process consisting of a priming step by GYG followed by an elongation carried out by GYS and then a branching step by GBE. **c**, Cryo-EM map, and model of the tetrameric GYS1:GYG1^ΔCD^ complex at 3.0 Å resolution. Individual GYS1 and GYG1 subunits are colored separately. **d**, Enlarged view of the GYG1 region interacting with GYS1. GYS1 is colored purple and GYG1 is colored coral. **e**, Residues Cys137, Cys189, and Cys251 form a cysteine rich pocket on GYS1 at the interface with GYG1. Inset: Different contour levels for the cryo-EM density of Cys137 and Cys189 are shown.

Bulk glycogen synthesis is carried out by GYS, a retaining glycosyltransferase (GT) belonging to the GT3 superfamily. GYS catalyses the successive addition of α-1,4-linked glucose residues to the non-reducing end of a growing polysaccharide chain, using UDP-glucose (UDP-glc) as the glucose donor with the release of UDP^5^. In mammals, GYS is present as two isoforms, GYS1 and GYS2, sharing ~69% sequence identity^6^. GYS1 is expressed in most tissues including the muscle and brain^7^, while GYS2 is expressed only in the liver. Mammalian GYS is the rate-limiting enzyme in glycogen biosynthesis, and its activity is regulated post-translationally by two mechanisms: activation by the effector glucose-6-phosphate (Glc6P)^8,9^ and inhibition by reversible phosphorylation^10^.

Reversible phosphorylation of GYS is mediated by several Ser/Thr-directed protein kinases, occurs at multiple sites, and is hierarchal in that different sites contribute to GYS inhibition in a specific order and to varying degrees^11^. At least 9 *in vivo* phosphorylation sites have been identified at the N- and C-termini of mammalian GYS1, in which sites 2 (Ser8), 2a (Ser11), 3a (Ser641), and 3b (Ser645) are found to play more significant roles^12,13^. Dephosphorylation, performed by glycogen-associated phosphatases of type 1 (PP1), significantly alters GYS kinetic properties such as increased affinity for UDP-glc and sensitivity to the Glc6P activator^14^. Glc6P binds to an allosteric site equipped with an arginine cluster, overcomes phosphorylation-dependent inhibition, and increases the enzyme’s susceptibility to PP1-mediated dephosphorylation. These two regulatory mechanisms of mammalian GYS have been described by a three-state conformational model, comprising the Tense (T)/inhibited state where GYS is phosphorylated, Intermediate (I)/basal state when unphosphorylated, and Relaxed (R)/activated state when Glc6P is bound^15–18^.

The pleiotropic PP1 comprises a catalytic subunit (PP1c) and a regulatory subunit (PP1r), the latter targeting the phosphatase to specific targets. 7 glycogen-targeting PP1r (PPP1R3A – PPP1R3G), characterised by the presence of an RVSF motif for PP1c binding, a glycogen-binding motif VxNxxFEKxV and a putative GYS binding motif WxNxGxNYx(I/L), have been described^19–21^. Among them, subunit 3C (PPP1R3C; also known as protein targeting to glycogen, PTG) is ubiquitously expressed in the brain, liver, and heart, and its gene knockout indirectly reduces GYS activation^22^. As such, these PP1 regulatory subunits are often considered activators of GYS1, although direct interaction between these proteins has not been definitively shown. Nevertheless, PTG is thought to function as a scaffold for glycogen metabolic enzymes, such as GYS, glycogen phosphorylase, and phosphorylase kinase^22^.

GYS1 has emerged as a therapeutic target for several glycogen storage diseases (GSD), including GSD type II (Pompe disease)^23^, GSD type IV (Andersen disease and adult polyglucosan body disease)^24^ and Lafora disease^25^. The root of these disorders is the accumulation of aberrant or normal glycogen in affected tissues, due to defective glycogen synthesis or breakdown. Downregulating GYS1 activity to interfere with glycogen chain elongation therefore could present a therapeutic opportunity. Despite this, inhibitor development for GYS1 has not progressed rapidly^23,24^, in part due to a lack of structural data, beyond that from bacterial^28–30^, *S. cerevisiae^16^* and *C. elegans*^31^ GYS orthologues, to guide drug discovery efforts. In this study we used cryo-electron microscopy (cryo-EM) to determine the structure of phosphorylated human GYS1 in different functional states and characterized the interactions with its functional partners, namely glycogenin GYG1 and the PP1 regulatory subunit PTG.

## Results

### Structure of human GYS1 with a minimal interacting region of GYG1

Unlike *C. elegans* gsy-1 and yeast Gsy2p, producing large yields of recombinant soluble human GYS1 alone for structural studies has proven a challenge. However, co-expression with its binding partner, human GYG1, in an insect expression system has allowed for the isolation of this ~ 0.5 mDa complex as shown previously^29,30^. Using the same system, we co-expressed and purified the full-length GYS1:GYG1^FL^ complex (Extended Data Fig. 1b) but found it recalcitrant for crystallization. This was likely due to a combination of flexible regions along with heterogeneous phosphorylation and glucosylation of GYS1 and GYG1 respectively, as reported previously^29,30^ and determined by denaturing mass spectrometry (Extended Data Fig. 1f). As the complex is of sufficient size, cryo-EM was attempted but the GYS1:GYG1^FL^ complex was prone to aggregation and gave heterogenous particle sizes (Extended Data Fig. 1d).

Human GYG1 consists of an N-terminal catalytic domain, flexible linker, and a small C terminal GYS1-interacting domain (Fig. 1a). The crystal structure of full-length *C. elegans* gsy-1 in complex with the last 34 residues of gyg-1 demonstrated that this highly-conserved C-terminal region forms a helix-turn-helix motif sufficient for interaction with GYS1^31^. In our attempts to improve the complex for crystallization, we designed bi-cistronic constructs encoding untagged human GYS1 (aa 1-737) and His_6_-GST-tagged human GYG1 C-terminus (aa 264-350 or aa 294-350). Co-expression with GYG1 294-350 allowed for recovery of sufficient quantities of soluble GYS1 (Extended Data Fig. 1a). This construct (GYS1:GYG1^ΔCD^) is multiply phosphorylated as detected by intact mass spectrometry (Extended Data Fig.1f). Using a coupled spectrophotometric assay, this truncated complex had similar GT activity to the wild-type GYS1:GYG1^FL^ complex, and for both complexes activity was stimulated by Glc6P (Extended Data Fig. 1g). Despite considerable effort no crystals were obtained of GYS1:GYG1^ΔCD^, however it showed improved behaviour in cryo-EM grids presenting less aggregation than GYS1:GYG1^FL^. Individual particles with a distinctive box-like shape were easily discernible and initial 2D classification resulted in classes representative of a tetrameric particle (Extended Data Fig. 1d, e).

We determined a 3.0 Å structure of the phosphorylated GYS1:GYG1^ΔCD^ complex with *D2* symmetry applied (Fig. 1c, Extended Data Fig. 2). The cryo-EM map has a resolution range from 2.9 Å at the core to 3.9 Å at the periphery of the complex, allowing for modelling of residues 13-289, 293-629, 637-645 of GYS1 and residues 317-349 of GYG1. As expected, the complex adopts a rectangular box-shape with residues 317-349 of GYG1 at each corner of the GYS1 homo-tetramer (Fig. 1c, Extended Data Fig. 3a). Each GYS1 monomer consists of two Rossmann domains and a tetramerization domain, and interacts with GYG1 in a 1:1 ratio (Extended Data Fig. 3b). GYS1 assembles into a dimer of dimers with two major interfaces (Fig. 1c, Extended Data Fig. 3a): a tetrameric interface formed by tetramerization domains (A/D, B/C interfaces) and a dimeric/regulatory interface (C/D, A/B interfaces). The latter is contributed by the regulatory helix α24 from each subunit, harbouring conserved arginine clusters. In this state, each GYS1 active site, located at the cleft between the two Rossmann domains, is in a closed conformation due to additional inter-subunit contacts at a minor interface (B/D, A/C)^16,28^. Here, helix α2 of Rossmann domain 1 contacts helix α16 of the tetramerization domain of the neighbouring subunit via a salt bridge between Glu78 and Lys429 along with a hydrogen bond between Leu107 and Arg430 (Extended Data Fig. 3c).

The interactions of GYG1 with GYS1 are very similar to that found in the *C. elegans* crystal structure^31^ (Fig. 1d, Extended Data Fig. 3f). GYG1 uses a helix (αA)-turn-helix (αB) motif to interact with helices α4, α9, and α10 of hGYS1, with a combination of hydrogen bonds and hydrophobic interactions (Fig, 1d, Extended Data Fig. 3f). Looking at the hGYS1 region where hGYG1 interacts we observed a cysteine-rich pocket of residues Cys137, Cys189, and Cys251 near the last α-helix of hGYG1 (Fig. 1e). The distances between Cys137 and Cys189 (3.39 Å), and between Cys189 and Cys251 (4.18 Å) are within disulphide-bonding distance. Lower threshold values of the cryo-EM density indeed suggest a possible disulphide bond between Cys137 and Cys189 (Fig. 1e inset), however due to its ambiguity we modelled all three cysteine residues as reduced. Without GYG1 this cysteine-rich pocket of GYS1 would be solvent-exposed, thus GYG1 may stabilise this region by preventing aberrant disulphide formation. The lack of this cysteine-rich pocket (Cys137, Cys189, Cys251) in yeast Gsy2p (replaced by Val126, Pro177, Ser240) and *C. elegans* GYS1 (replaced by Cys154, Leu207, Thr269) may explain the unique requirement of co-expressing GYG1 to stabilise GYS1 in human (Supplementary Fig. 1). It is interesting to speculate that these cysteines may act as redox switch as similarly found in human brain glycogen phosphorylase^34^ which should be investigated in future studies.

### The structural basis of phosphorylation sensing

As-purified GYS1 was highly phosphorylated (Extended Data Fig. 1f), which is representative of the inhibited/T state and supported by the lack of GT activity in the absence of Glc6P (Extended Data Fig. 1g). However, GYS1 in this state adopts a similar conformation with the *C. elegans* gsy-1 and yeast Gsy2p basal/I state structures, with R.M.S.D. of 0.95 Å and 0.93 Å respectively (Extended Data Fig. 3d). In eukaryotic GYS, the N- and C- termini harbour several phosphorylation sites that mediate inhibition^12,13^ (Fig. 2a), and each phosphorylated site has been suggested to interact with specific conserved arginine residues present on a regulatory helix α24^16,28^. In our 3.0 Å map, density was present for modelling the N- and C- termini (Fig. 2b, Extended Data Fig. 4).

**Fig. 2.**
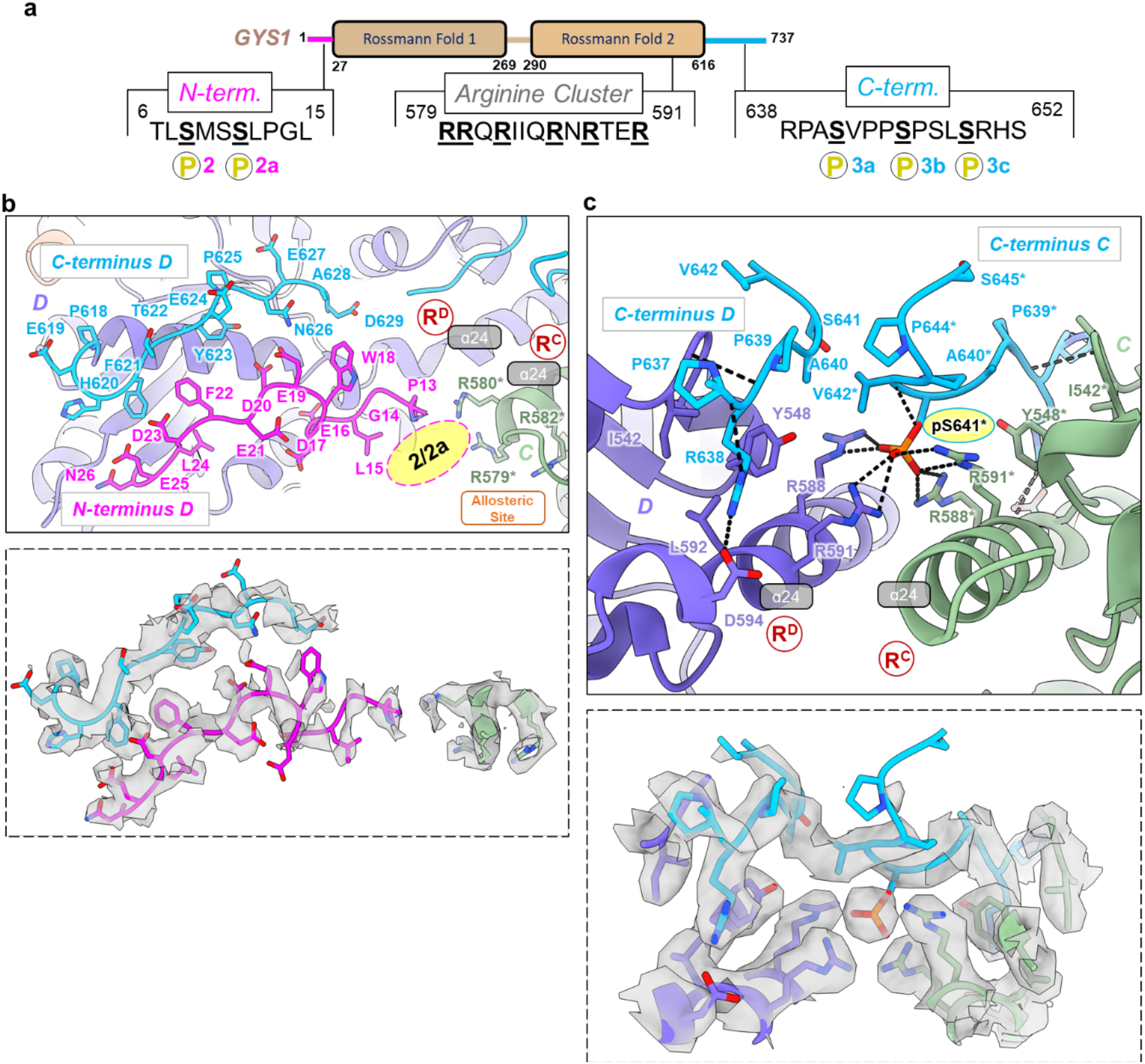
N- and C- termini of phosphorylated GYS1:GYG1^ΔCD^ complex in the inhibited/T state. **a**, Key sites of phosphorylation and arginine cluster of a GYS1 subunit. **b**, Model of the N- and C- termini from one subunit (D shown) pointing towards the allosteric sites and arginine clusters (R^C^, R^D^) at the dimeric C/D interface. Inset shows EM density of both termini along with arginine residues from the neighboring subunit that would interact with phosphorylation sites 2 and 2a. **c**, Model of the C-termini residues 637-645 from two neighboring subunits (C, D shown) interacting with their arginine clusters at the dimeric C/D interface. Inset shows EM density of both C-termini along with arginine clusters from both subunits interacting with a single site 3a phosphorylation (pS641). Asterisk indicates residues from the neighboring subunit. Arginine clusters containing helices α24 are labelled. Putative location for phosphorylation sites 2 and 2a are indicated by pink oval.

Both termini follow a trajectory different from that observed for the non-phosphorylated *C. elegans* gsy-1 basal/I state, and do not form any secondary structure (Extended Data Fig. 3e). In our inhibited/T state, the N- and C- termini from each subunit traverse from and towards the two regulatory helices α24 at the dimeric (C/D, A/B) interface, respectively. We modelled the N-terminus, from residue Pro13 onwards. Despite no clear density present for phosphorylation sites 2 (Ser8) and 2a (Ser11), based on our structure they are positioned near the regulatory helix α24 of the subunit across the dimeric interface, and close to both Arg579 and Arg580, which could potentially sense the phosphorylation at these sites (Fig. 2b). The N- and C-termini from one subunit traverse in antiparallel fashion towards its own regulatory helix α24, and the helix α24 from the subunit across the dimeric interface (Fig. 2b). Strong density was apparent, in both *C1* and *D2* symmetry maps, between Arg588 and Arg591 of both GYS1 subunits at the dimeric interface (Fig, 2c, Extended Data Fig. 4b). We can trace and model a single phosphorylated site 3a (Ser641) (Fig. 2c), which is the first C-terminus phosphorylation site in the sequence (Fig. 2a). The density of this region was symmetric in both the *C1* and *D2* symmetry maps (Extended Data Fig. 6c) and likely represents an average of different conformations of the C-termini. However, aided by both the unfiltered and LAFTER denoised maps (Extended Data Fig. 4c), C-terminal residues Pro637-Val642 for one subunit and Pro637-Ser641 for the other across the dimeric interface were modelled (Fig. 2c). This clearly shows that Arg588 and Arg591 from both subunits sense the phosphorylation from a single 3a site, at any time (Fig 2c). This implies that the other C-terminus from the dimeric interface is excluded by steric occlusion, and both C-termini appear to traverse away from the main body of the enzyme as evidenced by the density of the map (Extended Data Fig. 4d) and fuzzy protrusions from this region as seen in 2D classes (Extended Data Fig. 1e). Altogether, our model of this inhibited state suggests that the non-symmetric interaction of a single phosphorylated site 3a at the dimeric (C/D, A/B) interfaces, combined with inter-subunit interactions of phosphorylated sites 2/2a across the interface, stabilise the hGYS1 enzyme in the inhibited state.

### Allosteric activation by glucose-6-phosphate

To reveal GYS1 in the activated/R state, we determined a 3.7 Å resolution structure in the presence of the allosteric activator Glc6P (Fig. 3a, Extended Data Fig. 5), and a 3.0 Å resolution structure in the presence of both Glc6P and the glucose donor UDP-glc (Fig. 3b, Extended Data Fig. 6). Compared to the inhibited/T state, Glc6P induces large global structural changes in GYS1 that result in an outward rotation of ~35° of each subunit along the tetramer axis (Fig. 3a). This removes inter-subunit contacts at the minor interface (B/D, A/C) between the N-terminal Rossmann domain 1 of one subunit with the tetramerization domain of the neighbouring subunit (Extended Data Fig. 3c), freeing access to the active site between the Rossmann domains. When aligning one GYS1 subunit each from the inhibited/T and activated/R states, the tetramerization domain of the neighboring subunit (minor B/D, A/C interface) moves by ~18.6 Å away with respect to Rossman domain 1 (Fig. 3a, 4a). The increased flexibility of the N-terminal Rossman domain is quite evident in the EM map as this region is quite blurred and is of much lower resolution (~5.0 Å resolution) in comparison to the enzyme’s core (~3.6 Å resolution, Extended Data Fig. 5d).

**Fig. 3.**
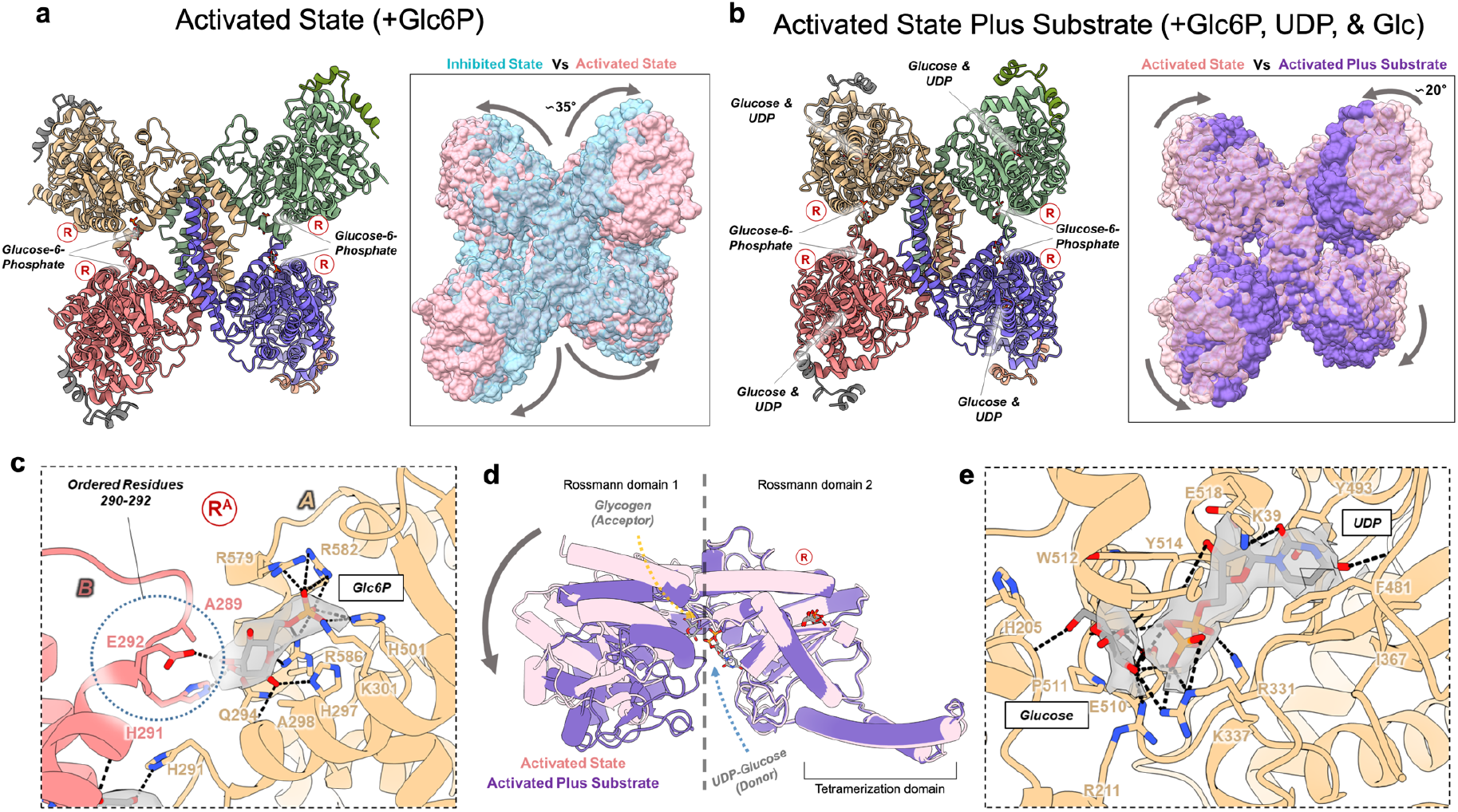
Activated structures of the phosphorylated R state GYS1:GYG1^ΔCD^ complex without and with substrate. **a**, Structure of the Glc6P bound activated (R) state determined from a 3.7 Å map. Inset shows the global conformational changes resulting from Glc6P activation in comparison to the inhibited (T) state. **b**, Structure of the activated (R) state bound to Glc6P, UDP, and glucose determined from a 3.0 Å map. Inset shows the global conformational changes resulting from substrate binding in the activated state. Regulatory/arginine cluster containing helices (α24) are labelled R. **c**, *Cis* and *trans* interactions with the Glc6P activator in the R state determined from the higher resolution substrate bound map. Interactions with glucose-6-phosphate in the lower resolution map without substrate are the same. Cryo-EM density for Glc6P is shown. **d**, Conformational changes of Rossmann domain 1 in relation to Rossmann domain 2 due to UDP and glucose binding in the R state. **e**, Interactions with UDP and glucose in the R state. Cryo-EM densities for both ligands are shown.

**Fig. 4.**
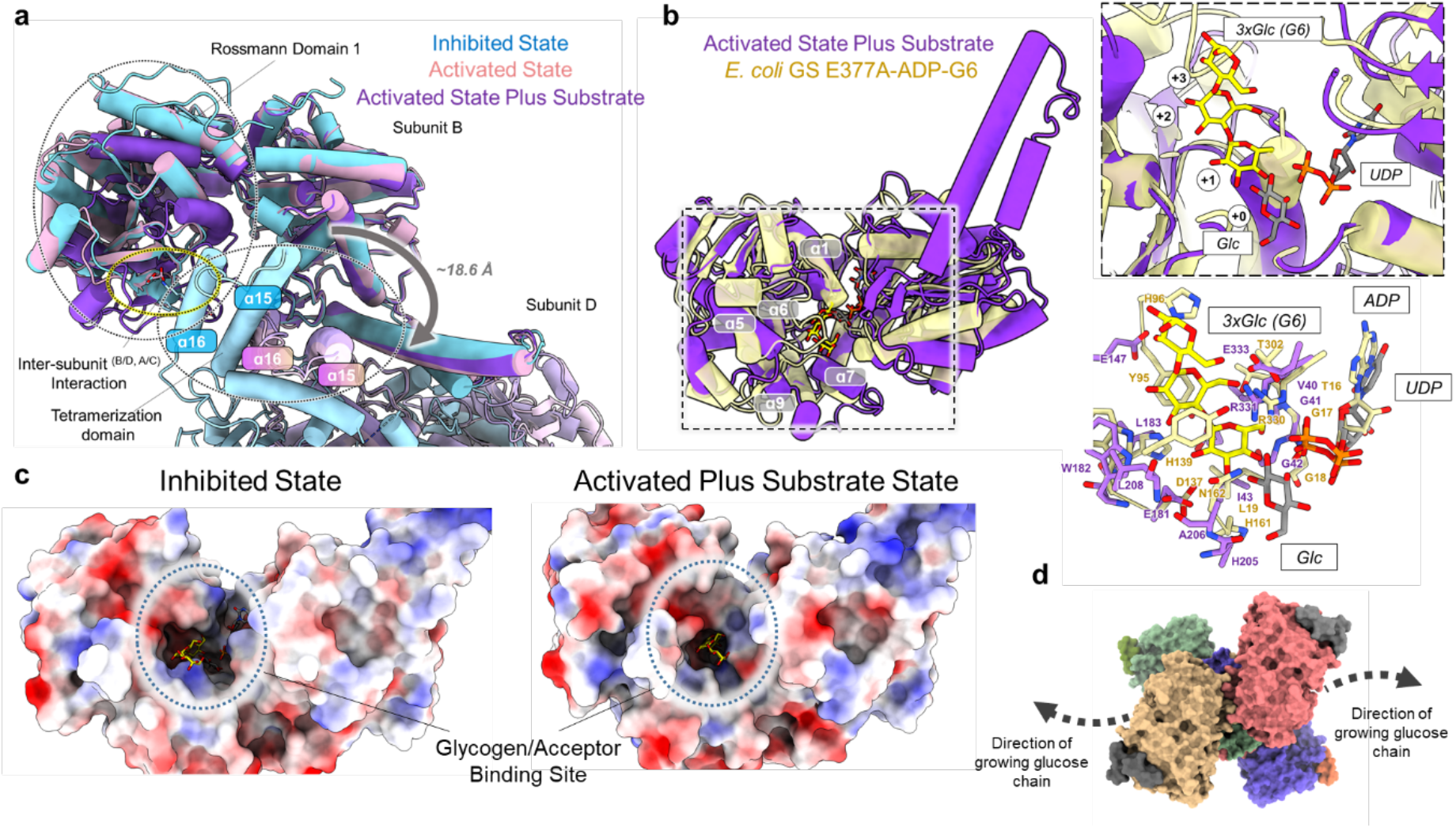
Structural comparison of the GYS1:GYG1^ΔCD^ inhibited and activated states with oligosaccharide bound *E. coli* glycogen synthase. **a**, Structural alignment of one subunit of the inhibited, activated, and activated plus substrate bound GYS1 structures. Tetramerization helices are highlighted to show relative movement between adjacent subunits within tetrameric hGYS1 **b**, Structural alignment of the activated plus substrate bound state against *E. coli* glycogen synthase incubated with maltohexaose (G6) bound with three glucose moieties in the active site. The first inset shows the active site of the two structures. The second inset demonstrates conservation of key residues involved in glucan binding **c**, Electrostatic surfaces of the inhibited and activated plus substrate bound states. The predicted glycogen binding site cleft is highlighted. **d**, Surface model of the activated state bound to UDP and glucose and the predicted direction of the growing glucose chain.

Glc6P binds identically to both R state structures, so we describe its binding mode based on the higher resolution structure bound with Glc6P and UDP-glc (Fig. 3c). Three arginines from the regulatory helix α24 (Arg579, Arg582, and Arg586), along with Lys301 and His501, interact with the phosphate moiety of Glc6P. The glucose moiety is recognised by His287, Gln294, and Arg586 from its own subunit (*i.e*. in *cis*), along with the now ordered residues (His291, Glu292) at the end of helix α13 from the neighbouring subunit across the dimeric interface (*i.e*. in *trans*). The binding mode of Glc6P and the disordered-to-ordered transition of residues 290-292 are conserved in the Glc6P bound yeast gsy2p crystal structure^16^. Ordering of this region is essential for the structural transition from the basal or inhibited state to the activated state (next section).

The activated/R state bound with UDP-glc is in a similar conformation to the activated/R state without UDP-glc (R.M.S.D. of 0.71 Å) except for a ~20° rotation of the Rossmann domain 1 relative to Rossmann domain 2, which closes the active site cleft (Fig. 3b, d). We observed density at the sugar donor site which fits better as individual UDP and glucose moieties, suggesting that UDP-glc was hydrolysed (Fig. 3e). This is similar to an activated structure of yeast gsy2p incubated with UDP-glc, in which one subunit has UDP and glucose bound^17^. Structural alignment shows that the Rossmann domain closure is identical to that of yeast gsy2p with (R.M.S.D. of 0.91 Å) and the UDP-glc binding residues are highly conserved (data not shown). In our structure, the uridine moiety of UDP is sandwiched between Ile367, Phe481, and Tyr493, also forming a hydrogen bond with Lys19 (Fig. 3e). The backbone of Gly41 and the sidechain of Glu518 interacts with the ribose moiety, while Arg331 and Lys337 disperse the charge of the diphosphate moiety. The hydrolysed glucose molecule forms multiple hydrogen bonds with the sidechains of Arg211, Arg311, Glu510, and Tyr514 along with the backbones of His205, Trp512, and Gly513. Additionally, Ala206 and Pro511 form hydrophobic interactions with the sugar (Fig. 3e).

This UDP-glc bound activated/R state is predicted to be the catalytic competent state poised for binding the glucose chain substrate^26,32^, and is different from the inhibited/T state as interactions of helix α2 with the central tetramerization domain at the minor interface (B/D, A/C) are still broken (Fig. 4a). The features of the N-terminal Rossmann domain 1 are also highly blurred (Extended Data Fig. 6d) suggestive of increased flexibility. To gain further insight into substrate binding and catalysis, we aligned one subunit of each of our states with the structure of *E. coli* glycogen synthase (GS) incubated with maltohexaose resulting in three glucose moieties bound to the active site (PDB <u>3CX4</u>)^35^. *E. coli* GS is in a closed conformation with respect to the active site and aligns with a R.M.S.D. of 1.09 Å and 1.19 Å against our hGYS1 inhibited and activated states respectively (Fig. 4b). We find the glucose moieties occupy the +1 to +3 sites while the hydrolyzed glucose in our EM map is in the +0 site (Fig. 4b). This predicted binding site of the glucan has conserved residues between *E. coli* GS and human GYS1 (Fig. 4b) and suggests that the initial growing glucose chain is fed into and then out of the GYS1 active site through a cleft formed by helices α1, α5, α6, α7, and α9 of Rossmann domain 1 (Fig. 4b, d). This pocket is not closed in the inhibited/T state and may explain the large increase in affinity for UDP-glc^36^ and glycogen when hGYS1 is in the activated/R state^37^ (Fig. 4c).

### Phosphorylation attenuates allosteric activation by glucose-6-phosphate

During the processing of the GYS1:GYG1^ΔCD^+Glc6P dataset, one 3D class appeared structurally similar to the inhibited/T state. This class was refined to 4.0 Å resolution with *D2* symmetry applied (Extended Data Fig. 7). While like our inhibited/T state map, where phosphorylated Ser641 of the C-terminus interacts with the arginine clusters, density for Glc6P in the allosteric site was apparent for this structure (Fig. 5a). Unlike the activated/R state, Glc6P in this structure does not interact with subunits in *trans* (across the dimeric interface) because residues 290-292, which interact with the glucose moiety in *trans* in the activated/R state, remain disordered. In this ‘inhibited-like’ state, all interactions involve the phosphate group and are identical to the activated states except for Arg586 which is not in a productive conformation to interact with both the glucose and phosphate moieties of Glc6P (Fig. 5c).

**Fig. 5.**
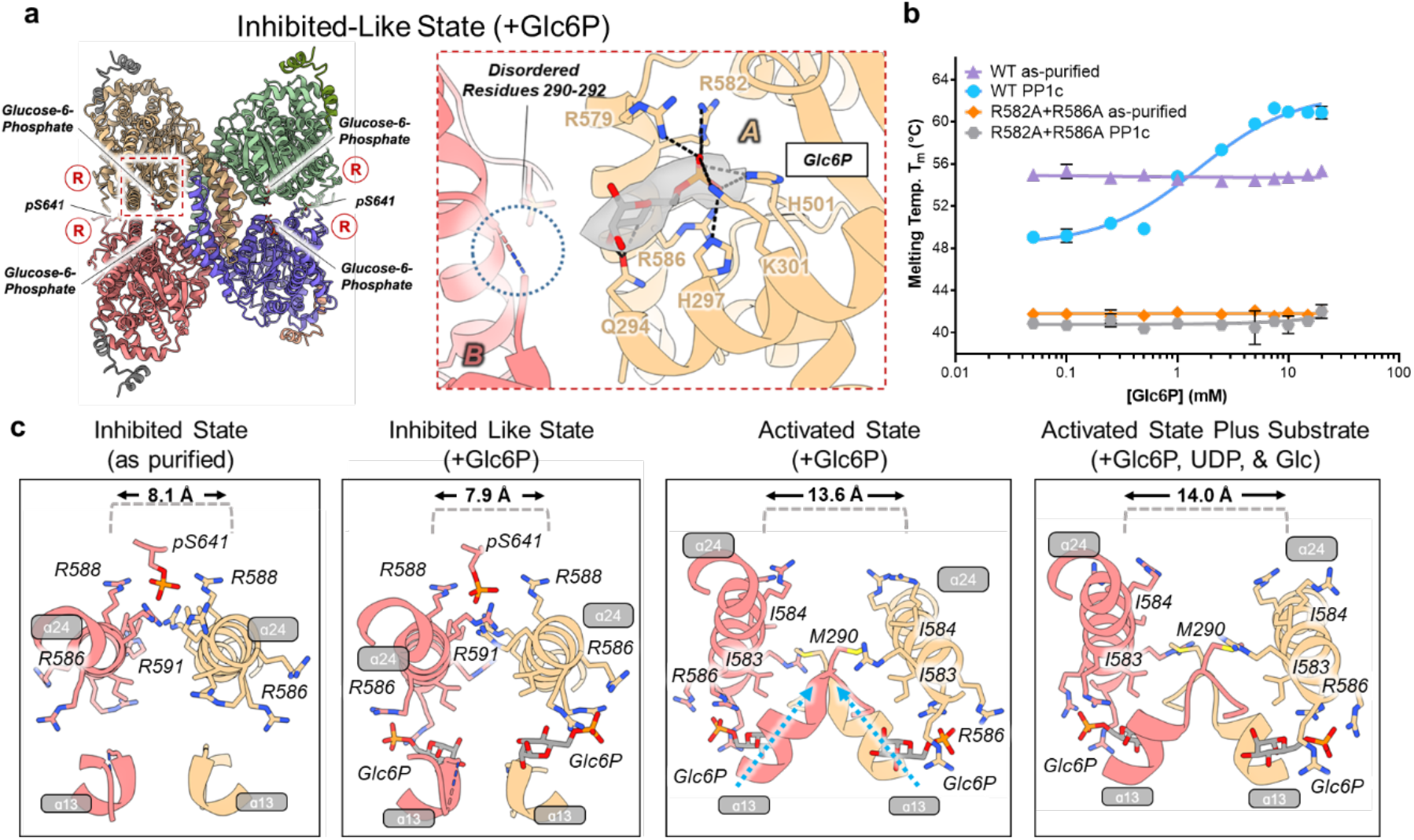
Phosphorylation hinders transition into the activated/R state as shown by the phosphorylated inhibited/T state bound to glucose-6-phosphate. **a**, Overall model of the phosphorylated T state bound to Glc6P and the interactions with this activator. Inset shows cryo-EM density for glucose-6-phosphate. Regulatory/arginine cluster containing helices (α24) are labelled R. **b**, Thermal shift assay of as-purified (phosphorylated) versus PP1c-treated (dephosphorylated) GYS1:GYG1^ΔCD^ (WT) and GYS1^p.R582A+p.R586A^:GYG1^ΔCD^ (R582A+R586A) complexes in the presence of increasing concentrations of glucose-6-phosphate. Median melting temperatures and standard deviations are shown (*n* = 4 technical repeats). **c**, R helix interactions and conformational changes as seen in our cryo-EM structures. Key residues are labelled. Distances between the R helices (α24) were determined as the distance between the Cα of the Asn587 residues.

With regards to allosteric activation, this ‘inhibited-like’ state potentially exists in dynamic equilibrium with the activated state. The binding of Glc6P is well known to overcome the inhibitory effects of phosphorylation, however reported k_a_ values of Glc6P for phosphorylated GYS1 vary between 0.33-1.8 mM from insect cell-expressed GYS1^29,30^ and between 0.8-1.9 mM for rabbit GYS1^38^. Dephosphorylation significantly reduces the amount of Glc6P to half maximally activate the enzyme (A_50_) within a range of ~3, ~10, or ~100 fold^39^. These diverse values likely reflect phosphorylation heterogeneity of each GYS1 sample and suggest an interplay between phosphorylation and Glc6P activation. To explore how this interplay impacts the complex at the molecular level, we applied the thermal shift assay and titrated Glc6P against our three complexes (GYS1:GYG1^FL^, GYS1:GYG1^ΔCD^, GYS1:GYG1^p.Y195F^), each in the as-purified (*i.e.* phosphorylated) and the PP1c-treated (*i.e*. shown to partially dephosphorylate the protein, particularly at key sites^19^) forms (Extended Data Fig. 1f). For all three complexes, dephosphorylation significantly reduced thermal stability by ~6°C (Fig. 5b, Extended Data Fig. 8), suggesting that the phosphorylated inhibited/T state is more stable than the dephosphorylated basal/I state. This is possibly due to the loss of stabilizing interactions of phosphorylated 2, 2a, and 3a sites with the arginine clusters. Significantly for all three constructs, Glc6P had no to little stabilizing effect towards phosphorylated complexes, whereas each dephosphorylated complex was readily stabilized by Glc6P with a maximal increase in melting temperature of ~8-12°C (Fig. 5b, Extended Data Fig. 8). The apparent AC_50_ (concentration of ligand to reach half maximal melting temperature) for each dephosphorylated construct was 1.7 ± 0.2 mM (GYS1:GYG1^FL^), 1.5 ± 0.2 mM (GYS1:GYG1^ΔCD^), and 0.9 ± 0.2 mM (GYS1:GYG1^p.Y195F^). These values are lower than the reported k_a_ values for dephosphorylated GYS1, likely due to differences in the remaining phosphorylation of the samples and/or pleiotropic effects from substrates^39^. Furthermore, a GYS1^p.R582A+p.R586A^:GYG1^ΔCD^ complex, in which two arginines that interact with Glc6P phosphate moiety were substituted, showed no stabilising effect from Glc6P when treated with the PP1c phosphatase, confirming their critical role in binding the allosteric activator (Fig. 5b).

Next, we compared the orientation of regulatory helices α24 among our four structures (Fig. 5c). From these, the ordering of residues 290-292 at the end of helix α13 (which interact with Glc6P in *trans* across the dimeric interface) appears to be the driver of conformational change from the inhibited/T to activated/R states. The ordering of these residues is associated with movement of helix α13 towards the regulatory helix α24 across the dimeric interface, positioning the hydrophobic Met290 (from α13) to interact with Ile583 and Ile584 (from α24). This drives apart the regulatory helices across the dimeric interface, distancing them from 8.1 Å to 13.6 Å and abolishes the ionic interactions of Arg588 and Arg591 from both subunits with the single phosphorylated Ser641. This replacement of ionic with hydrophobic interactions allows for greater flexibility between each subunit, as this distance increases further to 14.0 Å when the donor substrate is present (Fig. 5c).

To visualize this better, we applied 3D variability analysis to show that the activated/R states are far more flexible than the inhibited/T states (Extended Data Fig. 9a, Supplementary Videos 1-5). In both the activated/R states, the Rossmann domain 1 flexes onto Rossmann domain 2. This flexing movement is even more pronounced when UDP and glucose are bound to the active site. No such Rossmann domain closure is apparent in both inhibited/T states. However, 3D variability analysis for the Glc6P bound inhibited-like state showed a unique movement not observed in the inhibited state without Glc6P. This movement appears as a 2.0 Å expansion of the complex from the tetrameric interface (Supplementary Video 1, Extended Data Fig. 9b) and by flexibly fitting our inhibited state model, we observe that helix α13 moves towards the regulatory helices (Extended Data Fig. 9c, d). Such movement suggests that this inhibited-like state is primed to change into the activated state either by changes in dynamic equilibrium, binding of substrate and/or dephosphorylation by PP1. As we only incubated with 5 mM Glc6P for the phosphorylated GYS1:GYG1^ΔCD^ EM samples, these findings coupled with our thermal shift results suggest that the conformational change to the activated state is attenuated by the phosphorylation of site 3a and possibly 2/2a.

### Associated glycogen is the main driver of PTG recruitment to the GYS1:GYG1 complex

PP1 is the only phosphatase known that dephosphorylates GYS1 *in vivo* with assistance from a glycogen-targeting regulatory protein, such as PPP1R3C/PTG, which has been suggested to form a direct interaction with GYS1^19^. Attempts to express full-length human PTG were unsuccessful; we instead obtained soluble protein with a construct encompassing residues Leu134-Val259. This construct contains the carbohydrate binding module 21 (CBM21) domain (residues 149-257) in which the predicted glycogen binding motif VKNVSFEKKV (residues 175-184) and GYS binding motif WDNNDGQNYRI (residues 246-256) are present. Using the AlphaFold^40^ predicted model of the PTG(CBM21) domain, we overlayed two crystal structures of the starch binding domain from *R. oryzae* glucoamylase bound to maltotriose and maltotetraose at two different sites (starch binding sites I and II)^41^. The *R. oryzae* sites I and II align well with the GYS and glycogen binding motifs of the PTG(CBM21) respectively (Fig. 6a). Additionally, sequence alignment of all known glycogen-targeting PP1 regulatory subunits (PPP1R3 family) against the starch binding domain of *R. oryzae* glucoamylase showed that both VKNVSFEKKV and WDNNDGQNYRI motifs are highly conserved across all the CBM21 domains, suggesting that these two motifs in PTG(CBM21) are involved in glycogen binding (Supplementary Fig. 2), and implying that PTG(CBM21) does not form a direct interaction with GYS1.

**Fig. 6.**
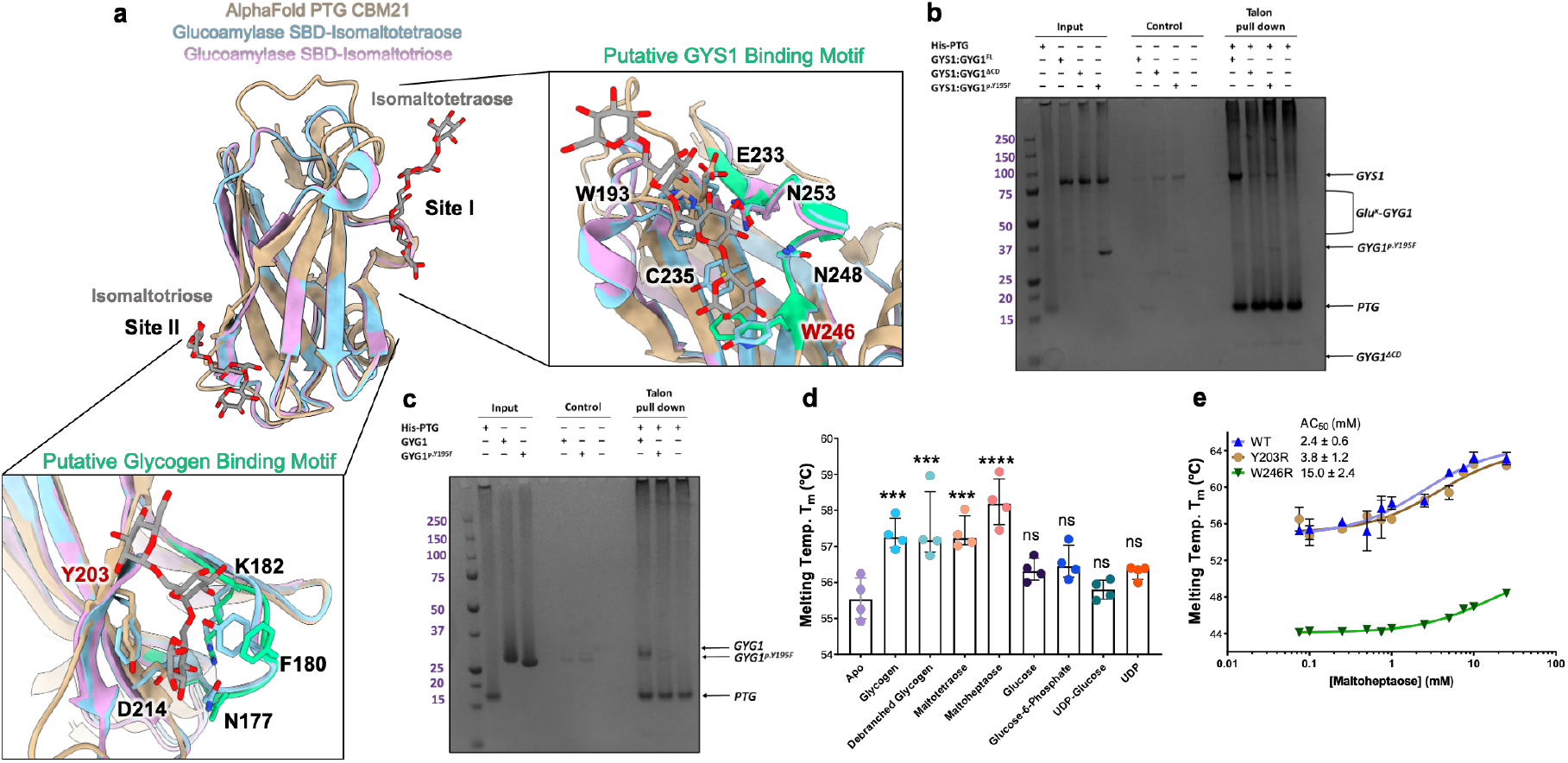
The CBM21 domain of PTG binds to the GYS1:GYG1 complex via the associated glucose chain. **a**, Structural alignment of the AlphaFold predicted structure of the PTG CBM21 domain against the starch binding domain from *R. oryzae* glucoamylase bound to maltotetraose and maltotriose at site I and site II respectively. Panels show how site I and site II align with the putative GYS1 binding motif and putative glycogen binding motif. Both motifs are coloured green. Y203 and W246 labels are highlighted red. **b**, PTG(CBM21) was incubated with GYS1:GYG1^FL^, GYS1:GYG1^p.Y195F^, or GYS1:GYG1^ΔCD^. The ability of PTG to bind GYS1:GYG1 complexes was assessed by affinity pulldown, followed by SDS–PAGE (*n* = 4 technical repeats). **c**, PTG(CBM21) was incubated with GYG1, or GYG1^p.Y195F^ catalytic domain constructs, passed onto affinity resin and analysed by SDS-PAGE (*n* = 4 technical repeats). GYG1 catalytic domain exists as a mixture of glucosylated states and runs at a higher apparent MW in SDS-PAGE than GYG1^p.Y195F^ which is non-glucosylated. **d**, Thermal shift analysis of PTG(CBM21) in the presence of various sugars and ligands (*n* = 4 technical repeats). Significance was determined by two-tailed unpaired t test. *** p<0.001; **** p<0.0001, ns: not significant. **e**, Thermal shift analysis of PTG(CBM21) wild-type (WT), PTG(CBM21)^p.Y203R^ variant (Y203R) and PTG(CBM21)^p.W246R^ variant (W246R) in the presence of increasing concentrations of maltoheptaose. Median melting temperatures and standard deviations are shown (*n* = 3 technical repeats).

To further characterize this, we used affinity pull-down to evaluate the binding of PTG(CBM21) to complexes of GYS1 and GYG1 (Fig. 6). His-tagged PTG(CBM21) pulled down only GYS1:GYG1^FL^ where GYG1 is attached with a glucose chain (glucosylated), at a level above background, but did not pull down GYS1:GYG1^ΔCD^ or GYS1:GYG1^p.Y195F^ complexes where GYG1 is not glucosylated (Fig. 6a and Supplementary Fig. 3a). This result agrees with analysis by blue-native PAGE (Supplementary Fig. 3c), suggesting that PTG(CBM21) is recruited to GYS1 by the GYG1-associated glycogen. To confirm a direct interaction between PTG and the GYG1 glucose chain, we repeated the PTG pull-down with the catalytic domain alone from GYG1 wild-type (glucosylated) and GYG1^p.Y195F^ (non-glucosylated), without GYS1. His-tagged PTG(CBM21) pulled down only glucosylated GYG1 catalytic domain at a level above background, but not the non-glucosylated GYG1^p.Y195F^ catalytic domain (Fig. 6b and Supplementary Fig. 3b).

Next, the polysaccharide binding ability of PTG(CBM21) was studied by thermal shift assay. Only glycogen, debranched glycogen, maltotetraose, and maltoheptaose resulted in increased stability of PTG(CBM21) (Fig. 6d). To demonstrate that the two sequence motifs of PTG(CBM21) are involved in sugar binding, we substituted to arginine the residues Tyr203 and Trp246, representing a conserved residue from the putative glycogen binding motif (equivalent to *R. oryzae* site II) and GYS binding motif (*R. oryzae* site I) respectively (Fig. 6a). Whereas PTG(CBM21)^p.Y203R^ had a similar melting temperature as the wild type PTG(CBM21), PTG(CBM21)^p.W246R^ was approximately 10 °C less stable (Fig. 6e). Titrating maltoheptaose stabilized both wild type PTG(CBM21) and PTG(CBM21)^p.Y203R^ similarly, with AC_50_ values of 2.4 ± 0.6 mM and 3.8 ± 1.2 mM respectively. In contrast the other variant PTG(CBM21)^p.W246R^ had a severely reduced ability to bind maltoheptaose, with an apparent AC_50_ of 15.0 ± 2.4 mM (Fig. 6e), showing that site I has a significant role in sugar binding. Overall, these results suggest that the GYG1-associated glycogen of a GYS1:GYG1 complex is the major binding site of PTG, and that any direct GYS1-PTG interactions are potentially quite weak, or outside of the domain boundaries of the CBM21, or only form in the presence of PP1.

## Discussion

Glycogen synthase is a metabolic enzyme underpinning the classic paradigms of protein allostery and phosphorylation-dependent regulation. Despite the well-characterised enzyme kinetics, and the discovery nearly a decade ago that recombinant GYS1 can be co-expressed with GYG1, until now structural information for human GYS1 has remained elusive. Taking advantage of a minimal GYG1 interacting polypeptide that introduces less disorder to the complex with GYS1, and the capability of cryo-EM to classify subtle protein conformational features, we have determined the structure of phosphorylated human GYS1 under several inhibited and activated states, allowing us to chart its trajectory between phosphorylation mediated inhibition and allosteric activation.

Our inhibited state structures have unravelled the roles of phosphorylated N- and C-termini as a molecular “straitjacket”, reducing the flexibility of the GYS1 tetramer and hindering the Glc6P-mediated conformational change to the activated state. Specifically phosphorylated site 3a, and potentially also sites 2/2a, are poised to interact with conserved arginine clusters at the dimeric interface, confirming their significance relative to other sites^39^. Sites 2/2a could interact with Arg579 and Arg580 in *trans* (subunit across the dimer interface). Unexpectedly we found that one single phosphorylation at site 3a interacts with Arg588 and Arg591 from both subunits at the dimeric interface (i.e. in both *cis* and *trans*). The essentiality of Arg579, Arg580, Arg588 and Arg591 for phosphorylation-dependent inhibition is supported by mutagenesis of equivalent residues in yeast gsy2p^16^ and mouse GYS1^36,39,40^. This is further underscored by reciprocal mutagenesis of sites 2/2a and 3a in rabbit GYS1 that ablated inhibition by phosphorylation^12,44^ and/or improved sensitivity towards Glc6P activation^45^. The relative contributions of site 2/2a and site 3 in inducing phosphorylation-dependent inhibition remains unclear, and translating biochemical findings from yeast, mouse and rabbit orthologues to understanding the human enzyme may also be hindered by the variation in their N-terminus lengths and sequences^16,38,39^.

The Glc6P binding site, involving Arg579, Arg582, and Arg586 of the arginine cluster, is highly conserved between yeast and human^16^. Particularly, the importance of Arg582 and Arg586 is confirmed by their substitution in rabbit and yeast GYS which abolished Glc6P activation^16,36,39,40^ agreeing with our findings for human GYS1 (Fig. 5b). The Glc6P induced conformational change is also conserved in yeast gsy2p^16^, and our four human structures provide further clarity, showing that the ordering of residues Met290-Glu292 to interact with Glc6P in *trans* across the dimer interface drives the conformational change. This positions Met290 in-between the two regulatory helices α24 at the dimer interface, driving them apart with steric hinderance against Ile583 and Ile584 of the *trans* subunit. Therefore, Glc6P activation replaces the ionic interaction of phosphorylation with a hydrophobic interaction, allowing for greater flexibility between subunits and between the Rossmann domains from a single subunit that increase active site access. The equivalent residues of Met290, Ile583 and Ile584 in yeast (Phe299, Ile584, Asn585) and *C. elegans* (Leu308, Ile604, Ile605) suggest a shared mechanism for allosteric activation of glycogen synthase as a homo-tetramer.

Dephosphorylation of GYS1 by PP1 also relieves inhibition of GYS1 by removing the phosphorylation at sites 2/2a and 3a thus releasing the “straitjacket” effects of the N- and C-termini^29,30,36^. PP1 is recruited to its substrate proteins by different regulatory subunits, of which seven are known to recruit it to glycogen^46^. Among them, PTG is ubiquitously expressed^47^ and considered a therapeutic target for GSDs^22^. All known glycogen-recruiting regulatory subunits differ in length, but share a PP1 binding motif and a CBM21 domain^21^. The latter contains two putative binding sites^20^, namely site II corresponding to a glycogen-binding motif VxNxxFEKxV^19–21^, and site I WxNxGxNYx(I/L) suggested to be a GYS binding motif by work on the CBM21 domain of muscle-specific PPP1R3A (65.7% sequence similarity with PTG)^19,45^ (Supplementary Fig. 2). Our pulldown experiments suggest that PTG(CBM21) does not interact directly with GYS1, contrasting with a recent study involving PPP1R3A and the full-length GYS1:GYG1 complex which did not account for GYG1 self-glucosylation^32^. Instead our mutagenesis results mirror previous findings on the starch binding domain of *R. oryzae* glucoamylase, where mutating the equivalent residue (Tyr94 corresponding to Trp246 in PTG) in site I severely reduced the binding affinity for carbohydrate^49^. These findings suggest rather that PTG (and possibly other glycogen-targeting PP1 regulatory subunits) recruits PP1 to GYS1 via the GYG1-attached glucose chain. With multiple surface sites in addition to the active site of GYS1 for glycogen contacts^15^, the PTG-glycogen interaction therefore provides for GYS1 processivity, by facilitating PP1 recruitment to flexibly dephosphorylate^50^ the many sites on the N- and C- termini of GYS1. It is however possible that a GYS1 binding site is formed in the context of full-length PTG or in complex with PP1 and therefore further investigation is needed.

Together with interaction studies of the PP1 regulatory subunit PTG, our structural snapshots of GYS1 reveal a model of its regulation by both Glc6P and phosphorylation, explaining how their interplay alters the equilibrium of the various GYS1 states, further elaborating the lock-and-key hypothesis of these two effectors (Fig 7)^16,36^. This dynamic system likely allows for fine tuning of glycogen formation in response to upstream messengers such as insulin^14^. Furthermore, our structures provide novel opportunities in rational drug design of GYS1 inhibitors for treatment of GSDs. The validity of GYS1 as a target is supported by proof-of-concept *GYS1* knockout in cell and animal models^46,47^, and a safety profile is underscored by healthy individuals with reduced GYS1 enzyme activity^48,49^. Preventing dephosphorylation by targeting PTG and targeting the Glc6P allosteric site appear to be ideal starting points for inhibitor design. Indeed ATP has been suggested to be a competitive inhibitor of Glc6P and may trap GYS1 in an inhibited state^39^. Overall, our structural work elucidates decades of studies on the arginine clusters, key phosphorylation sites, and the conformational flexibility of GYS1.

**Fig. 7.**
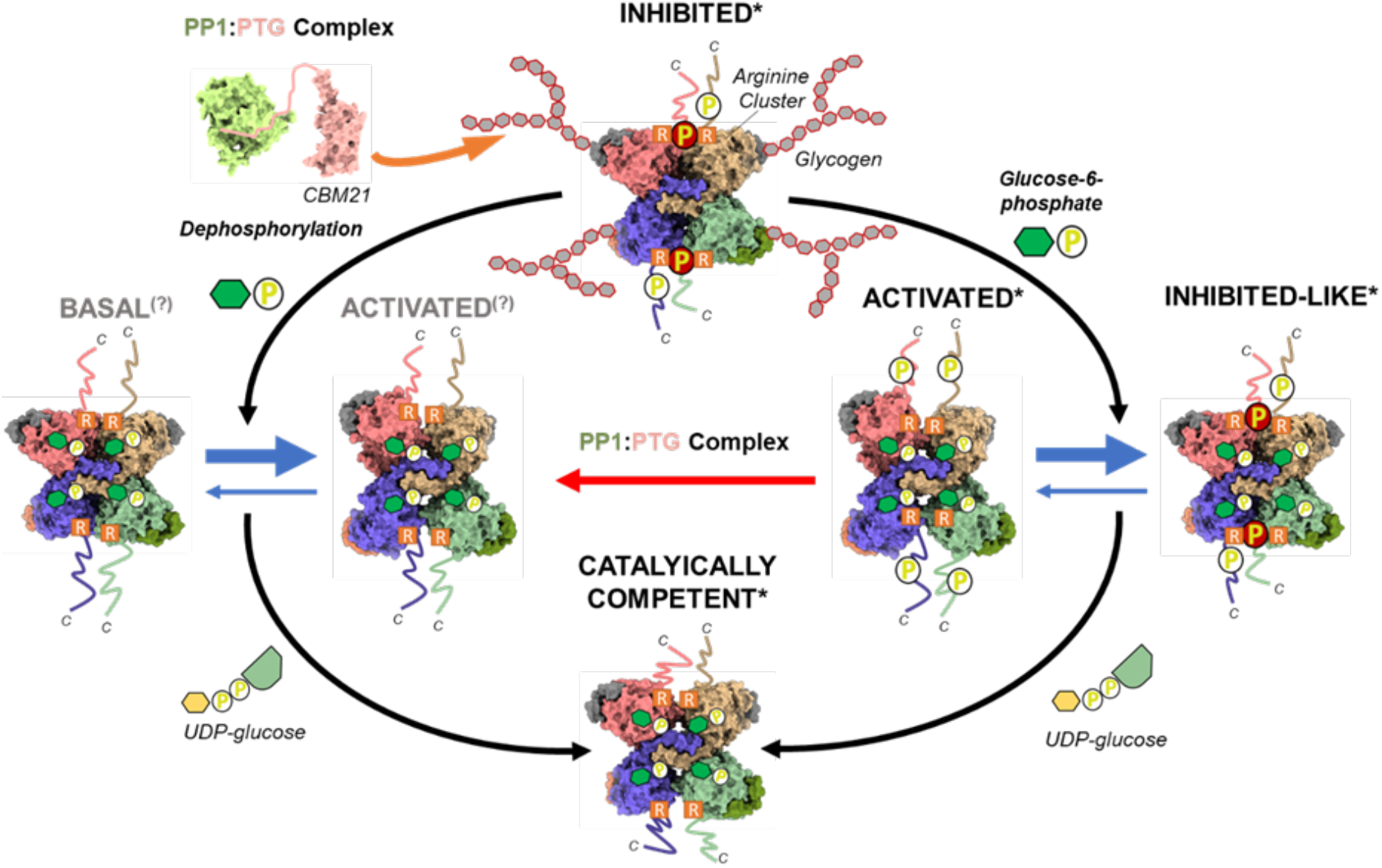
Proposed model of phosphorylation and glucose-6-phosphate regulation of hGYS1 activity. Only the C-termini and 3a phosphorylation site are shown for simplicity. Additionally, the associated glycogen is only shown for the inhibited state, though it is present in all other states. Structures with an asterisk are experimentally determined. Structures with a question mark are theoretical. Our model based on the structural data proposes that the inhibited/T state is catalytically inactive because the phosphorylated N- and C- termini bind to a subunit interface. This locking interaction reduces GYS1 flexibility and prevents active site closure by the two Rossmann domains. Glc6P binding to the allosteric site overcomes these inhibitory effects to promote a conformational change to the activated/R state. However, this activated/R state is in a dynamic equilibrium with an inhibited-like state, due to the competition between the locking interactions of phosphorylated termini at the subunit interface and the conformational change due to Glc6P binding. The inhibition of phosphorylation can also be relieved by the concerted actions of the PP1:PTG complex that binds to the associated glycogen and dephosphorylates the GYS1 N- and C- termini, resulting in the basal/I state. This intermediate state is more susceptible to the allosteric effects of Glc6P binding, shifting the dynamic equilibrium more towards the activated state. In the activated state binding of the substrate UDP-glc promotes the closure of the cleft between the two Rossman domains resulting in a catalytically competent state for extending the associated glycogen chain.

**Table 1.**
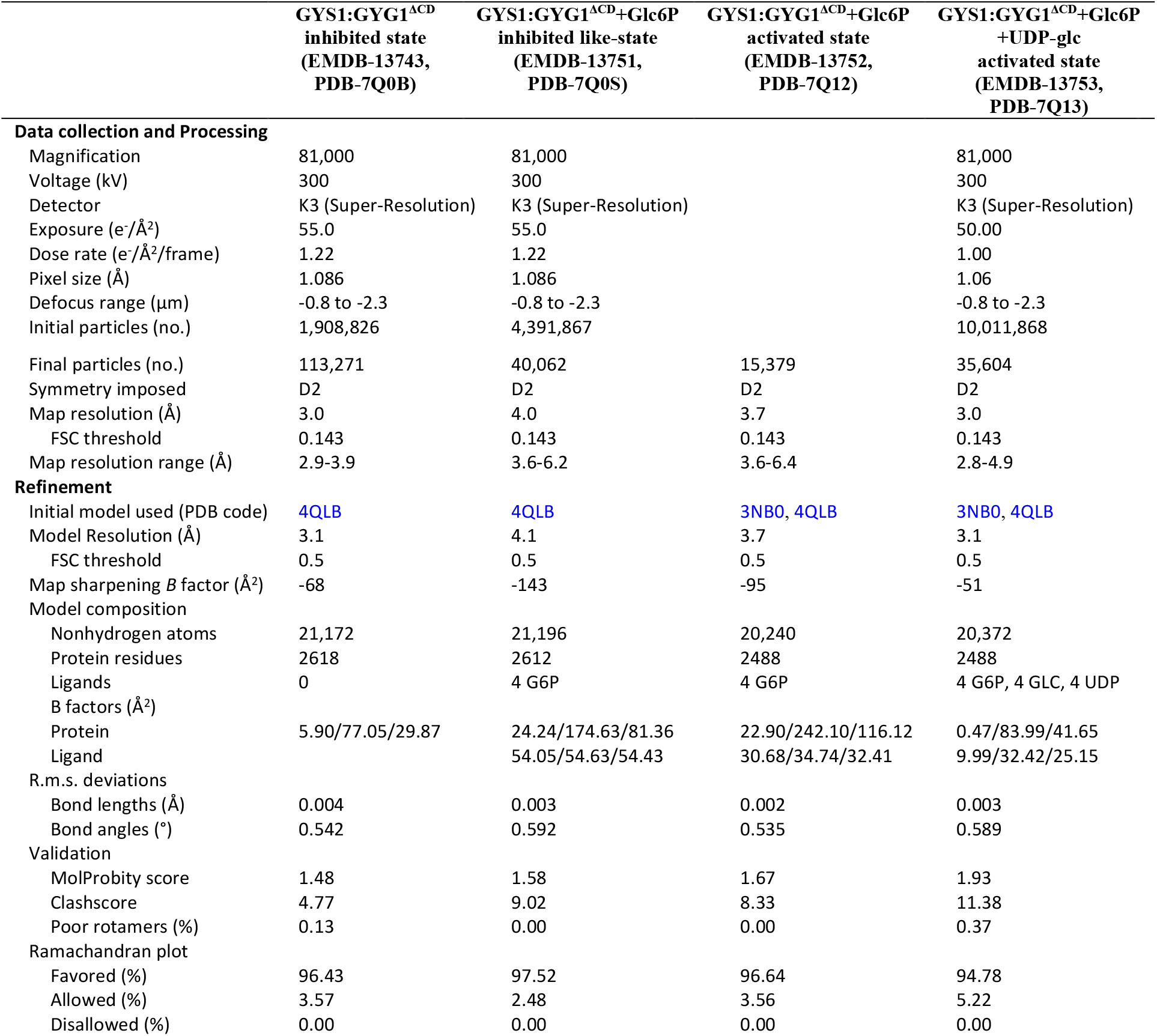
Cryo-EM data collection, refinement, and validation statistics

## Supporting information

SupplementaryVideo1_Component1

SupplementaryVideo1_Component2

SupplementaryVideo1_Component3

SupplementaryVideo1_Component4

SupplementaryVideo1_Component5

## Methods

### Cloning, expression, and purification of GYS1:GYG1 complexes

DNA encoding the full-length genes of human GYS1 (IMAGE: 3143019) and GYG1 (IMAGE: 3504538; isoform GN-1L with UniProt ID P46976-1) were amplified from a cDNA clone and subcloned into the FastBac™-Dual vector (Life Technologies) with an N-terminal His_6_-tag and a TEV protease cleavage site on GYS1. The GYG1^p.Y195F^ mutant was generated from this plasmid using the QuickChange mutagenesis kit (Stratagene). Codon optimised genes for GYS1 and aa 264/294-350 GYG1 (GYG1^ΔCD^) (with a stop codon) interspersed with a SV40 terminator and a polyhedrin promotor were artificially synthesised (Twist Biosciences). Codon optimised sequences for either a N-terminal TEV cleavable MBP-His_6_, His_6_-GST, or His_6_-GFP tag was appended to the GYG1 gene to allow purification. The resulting bistronic fragment was then inserted into pFB-CT10HF-LIC for insect cell expression. In-Fusion HD (Takara) mutagenesis was used to introduce specific mutants in the coding sequence of GYS1. All GYS1:GYG1 complexes were expressed in Sf9 cells grown in Sf-900™ III SFM (Life Technologies). Cell pellets were harvested, homogenized in lysis buffer (50 mM sodium phosphate pH 7.5, 500 mM NaCl, 5% glycerol, 0.5 mM TCEP, 10 mM imidazole) and insoluble material was removed by centrifugation. The GYS1:GYG1 complexes were purified by affinity (Ni-Sepharose; GE Healthcare) and size-exclusion (Superose 6; GE Healthcare) chromatography. Protein was treated with His-tagged TEV protease overnight at 4 °C, and then passed over Ni-Sepharose resin to remove the TEV protease and uncleaved protein. Purified complexes were concentrated to 10-20 mg/mL and stored in storage buffer (25 mM HEPES pH 7.5, 500 mM NaCl, 5% glycerol, 0.5 mM TCEP) at −80 °C.

### Cryo-EM sample preparation and data acquisition

GYS1:GYG1^ΔCD^ was diluted to 0.75 mg/ml into 25 mM HEPES, pH 7.5, 200 mM NaCl, 2.0 mM TCEP, 0.05% (v/v) tween-20 for the as purified, inhibited state. For the activated states of GYS1:GYG1^ΔCD^ was diluted to 0.75 or 0.5 mg/ml into 25 mM HEPES, pH 7.5, 200 mM NaCl, 2.0 mM TCEP, 0.05% (v/v) tween-20, 5 mM Glc6P, and 5 mM UDP-Glc when stated. Grids were prepared using a FEI Vitrobot Mark III at 4 °C and 100% humidity. 3 μl of sample was applied to a plasma treated gold coated R 1.2/1.3 300 mesh holey carbon grid (Quantifoil), with a blot force of 0, a blot time of 3 seconds and a wait time of 10 seconds.

Movies of GYS1:GYG1^ΔCD^ as purified and in the presence of Glc6P were collected during the same session at the Midlands Regional CryoEM Facility on a FEI Titan Krios equipped with a K3 (Gatan) direct electron detector operating in super-resolution mode. Images were imaged at 300 kV with a magnification of 81,000×, corresponding to a physical pixel size of 1.086 Å (super resolution pixel size of 0.543 Å). 45 frames over 5 seconds were recorded with a defocus range of −0.8 μm to - 2.3 μm with a total dose of (1.22 e^−^ A^−2^ per frame). Movies of GYS1+GYG1^ΔCD^ in the presence of Glc6P and UDP-Glc were collected at eBIC (Diamond Light Source) on a FEI Titan Krios equipped with a K3 (Gatan) direct electron detector operating in super-resolution mode. Images were imaged at 300 kV with a magnification of 81,000×, corresponding to a physical pixel size of 1.06 Å (super resolution pixel size of 0.53 Å). 50 frames over 3.4 seconds were recorded with a defocus range of - 0.8 μm to −2.3 μm with a total dose of (1.00 e^−^ A^−2^ per frame).

All datasets were corrected for beam induced motion with MotionCor2^53^ and CTF was estimated using CTFFIND-4.1^54^. Particles were auto-picked using the Relion 3.1.1^55^. Laplacian of Gaussian function and all further processing was done in Relion 3.1.1. For more detailed information on the processing workflow for all datasets please see extended data figures 2, 5, 6, and 7. All final maps were automatically sharpened in Relion 3.1.1. and for all but the inhibited state, locally filtered by resolution using LocRes. LAFTER^56^ maps were produced in aid of model building. Relion extracted particles and maps were imported into CryoSPARC v3.1.0 to use for 3D variability analysis^57^ with five components. Components were visualized by 3DVA simple display with 20 frames each using UCSF Chimera.

### Model fitting, refinement, and validation

Initial models of human GYS1 and GYG1 were built using the SWISS-MODEL server^55^ with structures of the *C. elegans* GYS1:GYG1^ΔCD^ and the activated Glc6P bound state of yeast Gsy2p (PDB <u>4QLB</u> and <u>3NB0</u> respectively) as a template. GYS1 models were docked into maps using Molrep^59^ and GYG1 was manually docked using UCSF Chimera. For the GYS1:GYG1 ^ΔCD^+Glc6P+UDP-glc activated state map Namdinator^60^ was used to flexibly fit the refined GYS1:GYG1^ΔCD^ + Glc6P activated model. Further model building and manual refinement was performed in COOT^61^ followed by iterative cycles of real-space refinement performed in Phenix^59^. Final models were validated using MolProbity^63^. Figures were created in UCSF Chimera and Chimera X^64^.

### Dephosphorylation of GYS1:GYG1 complexes

GYS1:GYG1 complexes at 5.0 mg/ml were dephosphorylated with 0.5 mg/ml PP1c in 25 mM HEPES, pH 7.5, 200 mM NaCl, 2.0 mM TCEP, 2.0 mM MnCl2 at room temperature for one hour. Reactions were halted by putting them into ice.

### UDP-Glo activity assay

The activity of GYS1:GYG1 complexes was measured using the UDP-Glo™ glycosyltransferase assay (Promega) according to manufacturer’s instructions. To measure activity 10 μL/well of reaction containing 100 nM GYS1:GYG1, 1 mM UDP-glucose, 0.5 mg/ml glycogen, and 10 mM glucose-6-phosphate in assay buffer (25 mM HEPES, pH 7.5, 200 mM NaCl, 0.5 mM TCEP) were dispensed into 384-well assay plates (Greiner). Following a 60-minute incubation at room temperature, 10 μL of UDP-Glo Plus detection reagent was added (final assay volume: 20 μL/well), and after a further 60 minutes room temperature incubation, luminescence was detected using a SpectraMax M3 (Molecular Devices).

### Cloning, expression, and purification of GYG1 and PTG

GYG1 was purified as previously described^62^. Human PTG (PPP1R3C) aa 134-259 (IMAGE clone: 4245774) was subcloned into the pNIC28-Bsa4 vector (GenBank accession no. EF198106) incorporating an N-terminal TEV-cleavable His_6_-tag. In-Fusion HD (Takara) mutagenesis was used to introduce specific mutants in the coding sequence of PTG. PTG was cultured in auto-induction Terrific Broth (Formedium) at 37 °C and induced overnight at 18 °C. Cell pellets were harvested, homogenized in lysis buffer (50 mM sodium phosphate pH 7.5, 500 mM NaCl, 5% glycerol, 0.5 mM TCEP, 10 mM imidazole) and insoluble material was removed by centrifugation. The supernatant was purified by affinity (Ni-Sepharose; GE Healthcare) and size-exclusion (Superdex 75; GE Healthcare) chromatography. Purified protein was concentrated to 10-20 mg/mL and stored in storage buffer (25 mM HEPES pH 7.5, 500 mM NaCl, 5% glycerol, 0.5 mM TCEP) at −80 °C.

### Talon pull down assay

His-PPP1R3C (1.0 mg/ml) was pre-incubated with either GYS1:GYG1 complex (0.25 mg/ml) or GYG1 (0.5 mg/ml) for 30 min at 4 °C in a total volume of 100 μl. Next 80 μl of a 50% slurry of Talon resin (Clontech) in binding buffer (25 mM HEPES, pH 7.5, 100 mM NaCl, 1 mM TCEP, 0.2% Tween 20) was added and incubated for a further 30 min at 4 °C. The resin was washed with 2 ml binding buffer with 10 mM imidazole and eluted with 40 μl 4× SDS PAGE sample buffer. Samples were run on SDS-PAGE and stained with Coomassie blue.

### Thermal shift assay

His-PPP1R3C or GYS1:GYG1 complex was diluted in thermal shift buffer (25 mM HEPES, pH 7.5, 200 mM NaCl, 2.0 mM TCEP) to 0.1 mg/ml with SYPRO-Orange (Invitrogen) diluted 1000X and with ligand at 1 mM in a total volume of 20 μl. Protein with ligand was incubated for 5 min at room temperature in 96-well PCR plates, before the addition of SYPRO-Orange. A Mx3005p RT-PCR machine (Stratagene) with excitation and emission filters of 492 and 610 nm, respectively was used to measure temperature shifts. AC_50_ values (half-maximal effective ligand concentration) were determined by fitting the melting temperatures using GraphPad Prism (v.9; GraphPad Software).

### Blue-Native PAGE

Blue-NATIVE PAGE was carried out as previously described and according to manufacturer’s instructions (Life Technologies). His-PPP1R3C, GYS1:GYG1 complex, and/or GYG1 were diluted in thermal shift buffer (25 mM HEPES, pH 7.5, 200 mM NaCl, 1.0 mM TCEP) was preincubated for 5 min at room temperature. All blue-native PAGE experiments were performed thrice independently.

## Data availability

Structures and EM maps of GYS1:GYG1^ΔCD^ inhibited state (EMDB-13743, PDB-7Q0B), GYS1:GYG1 ^ΔCD^+Glc6P inhibited like-state (EMDB-13751, PDB-7Q0S), GYS1:GYG1 ^ΔCD^+Glc6P activated state (EMDB-13752, PDB-7Q12), and GYS1:GYG1^ΔCD^+Glc6P+UDP-Glc activated state (EMDB-13753, PDB-7Q13) have been deposited to the EMDB and PDB databases.

## Acknowledgements

We thank all members of the SGC Biotech team especially Dong Wang for her molecular biology support. We thank Sean Froese for GYG1 constructs cloned at the SGC. We thank the Oxford Particle Imaging Centre (OPIC) electron microscopy facility for initial grid screening and data collection along with Laura Díaz Sáez for her help in the initial EM screening and data collection. We acknowledge The Midlands Regional CryoEM Facility at the Leicester Institute of Structural and Chemical Biology (LISCB), major funding from MRC (MC_PC_17136). We specifically want to thank Christos Savva for assistance and guidance in collection of data at LISCB. We acknowledge the Diamond Light Source for access and support to the UK’s Electron Bio-imaging Centre (eBIC, under BAG proposal EM20223) funded by the Wellcome Trust, MRC, and BBRSC. We specifically want to thank Peter Harrison for assistance in collection of data at eBIC. We also wish to thank Brian Marsden and Chris Sluman for their bioinformatics support. T.J.M. received cryo-EM training through Wellcome/MRC funded program (218785/Z/19/Z). The Structural Genomics Consortium is a registered charity (Number 1097737) that receives funds from AbbVie, Bayer Pharma AG, Boehringer Ingelheim, Canada Foundation for Innovation, Eshelman Institute for Innovation, Genome Canada, Innovative Medicines Initiative (EU/EFPIA) [ULTRA-DD grant no. 115766], Janssen, Merck & Co., Novartis Pharma AG, Ontario Ministry of Economic Development and Innovation, Pfizer, São Paulo Research Foundation-FAPESP, Takeda, and Wellcome Trust [092809/ Z/10/Z]. I.M.F. was supported by CNPq, Brazilian National Council for Scientific and Technological Development. T.J.M. and W.W.Y. also gratefully received Emerging Science Funds from Pfizer Worldwide Research and Development.

## Author contributions

T.J.M., A.B., and W.W.Y. designed the experiments. T.J.M. designed the GYS1:GYG1^ΔCD^ constructs, carried out mutagenesis, expressed and purified all GYS1:GYG1 complexes and PTG constructs, screened and collected EM data, analyzed, and refined all GYS1:GYG1^ΔCD^ structures and carried out all biochemical experiments. I.M.F. and L.S. carried out initial GYS1:GYG1 construct cloning, testing and optimization. D.S.F. expressed and purified the GYG1 constructs. T.J.M. and W.W.Y. analyzed the data and wrote the manuscript with contributions from all authors

---

**Extended Data Fig. 1.**
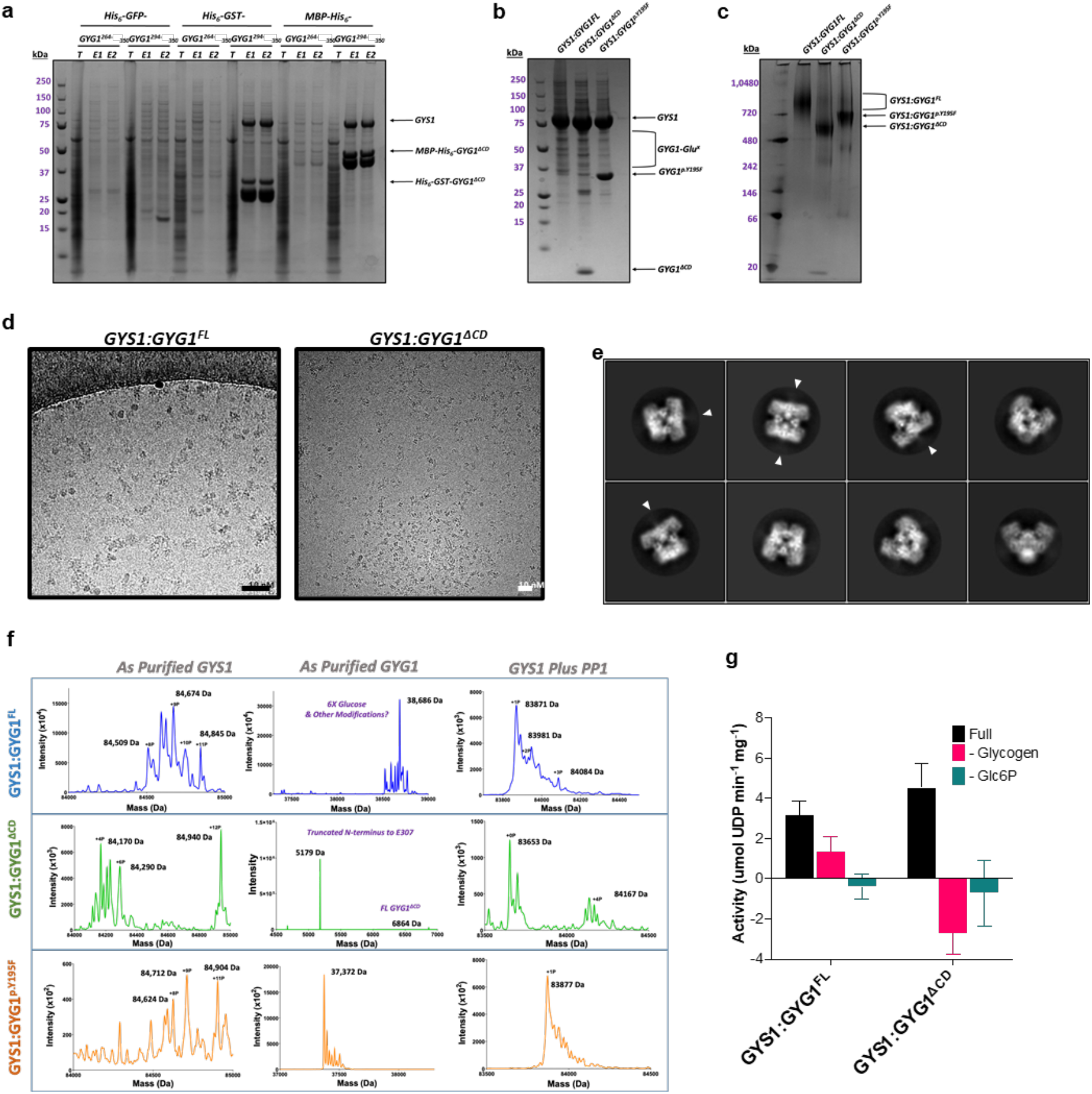
Purification and preliminary characterization of GYS1:GYG1 complexes. **a**, Coomassie stained SDS-PAGE of small-scale test purifications of GYS1 complexed with differently tagged truncated GYG1. **b**, Coomassie stained SDS-PAGE of the three GYS1:GYG1 complexes used in this study. **c**, Blue native PAGE of the three GYS1:GYG1 complexes used in this study. **d**, Example micrographs of GYS1:GYG1^FL^ and GYS1:GYG1^ΔCD^ complexes collected using a Glacios microscope. **e**, 2D classes of the GYS1:GYG1^ΔCD^ complex from an initial dataset collected using a Glacios microscope. Arrows indicate regions of fuzzy density protruding from an inter-subunit interface. **f**, Denaturing mass-spectra of GYS1 and GYG1, as purified and treated with PP1. **g**, UDP-Glo activity assay of the three GYS1:GYG1 constructs without and with exogenous glycogen. ‘Full’ is the activity assay with all substrates. ‘-Glycogen’ is the assay carried out in the absence of exogenously added glycogen. ‘-Glc6P’ is the assay carried out in the absence of Glc6P. Median and standard deviation of activity is shown (n = 3 technical repeats).

**Extended Data Fig. 2.**
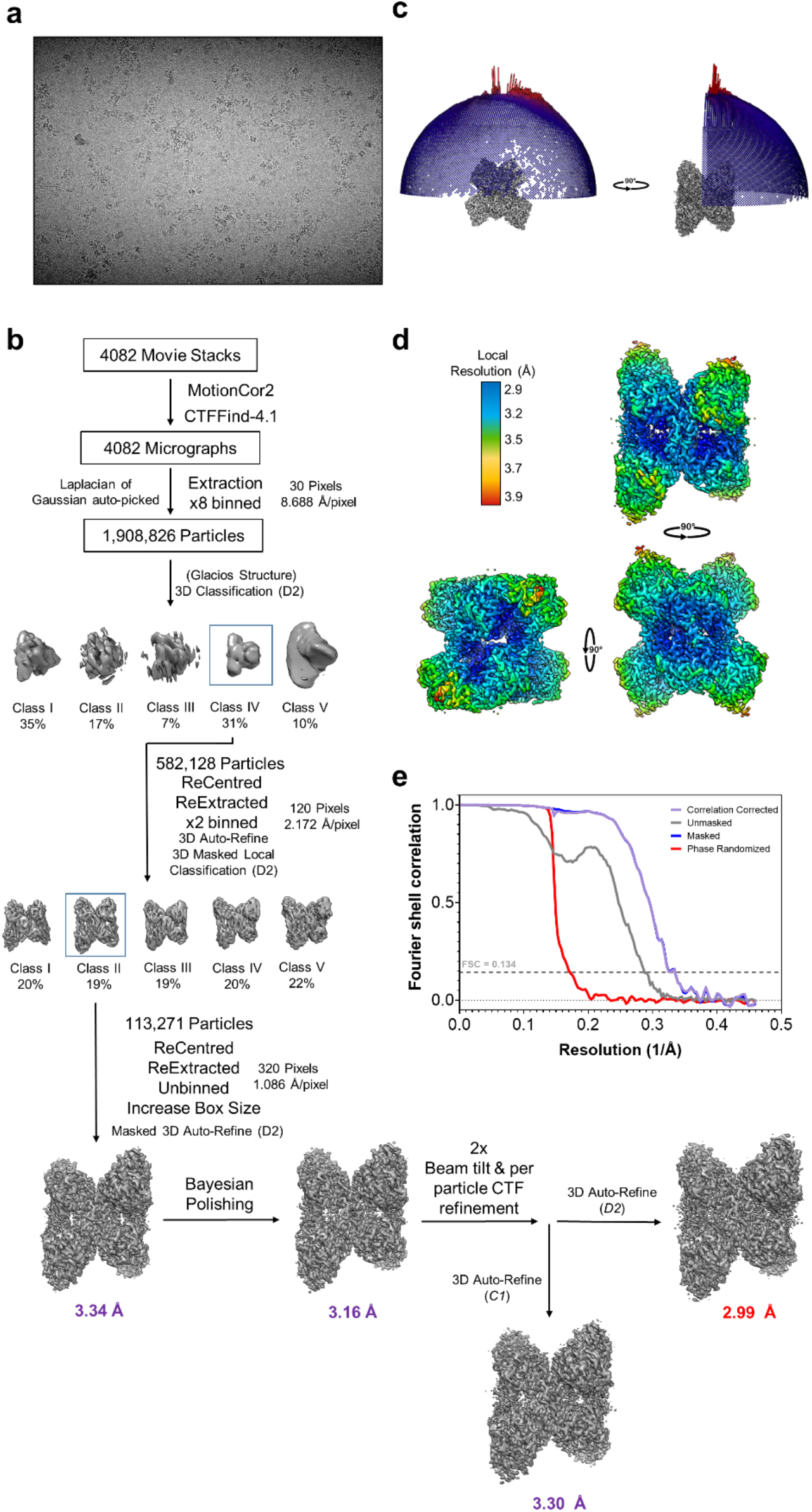
Image processing workflow of the GYS1:GYG1^ΔCD^ inhibited state. **a**, Representative K3 micrograph of the GYS1:GYG1^ΔCD^ inhibited state. **b**, Processing flow chart of the GYS1+GYG1^ΔCD^ inhibited state. **c**, Angular distribution of the 3.0 Å GYS1:GYG1^ΔCD^ inhibited state map. **d**, Local resolution variation of the 3.0 Å GYS1:GYG1^ΔCD^ inhibited state map. **e**, FSC curve of the 3.0 Å GYS1:GYG1^ΔCD^ inhibited state map.

**Extended Data Fig. 3.**
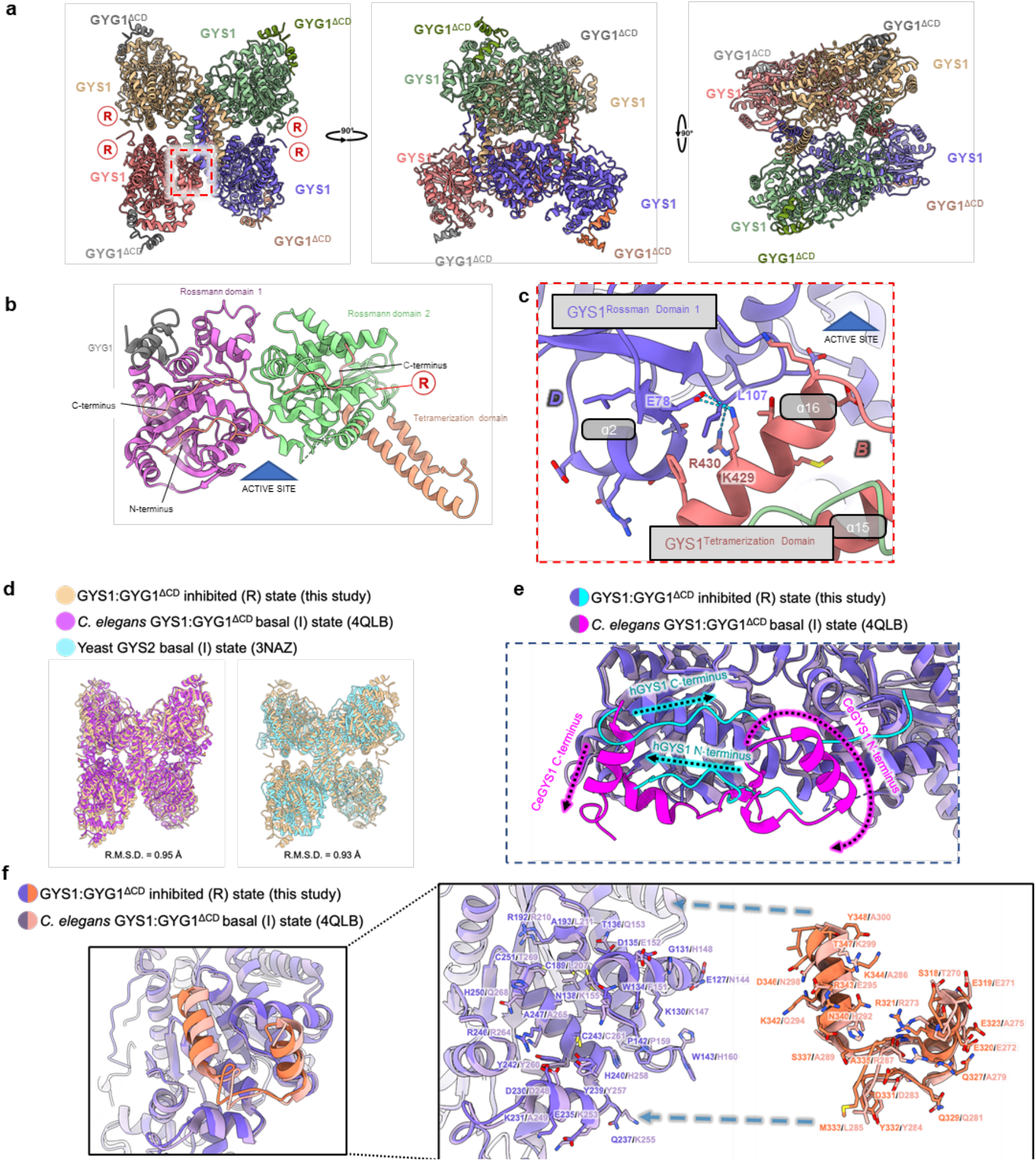
Structure of the GYS1:GYG1^ΔCD^ inhibited state and comparison with the *C. elegans* gys-1 and yeast Gsy2p basal/intermediate state structures. **a**, Model of the GYS1:GYG1^ΔCD^ inhibited state in three orthogonal views. R represents the location of the regulatory helix. **b**, Structural model of a GYS1:GYG1^ΔCD^ subunit showing the three domains of GYS1 as well as the GYG1 C-terminus. **c**, Close up of the inter-subunit interactions close to the active site cleft. **d**, Structural alignment of the inhibited/T state of the human GYS1:GYG1^ΔCD^ complex with the basal/I states of yeast gsy2p and *C. elegans* GYS1:GYG1^ΔCD^ complex. **e**, A zoom in view of the GYG1 interacting region of GYS1 of human and *C. elegans*. **f**, A structural alignment of the inhibited/T state of human GYS1 against the basal/I state of *C. elegans* GYS1 highlighting the different trajectories of the N- and C- termini.

**Extended Data Fig. 4.**
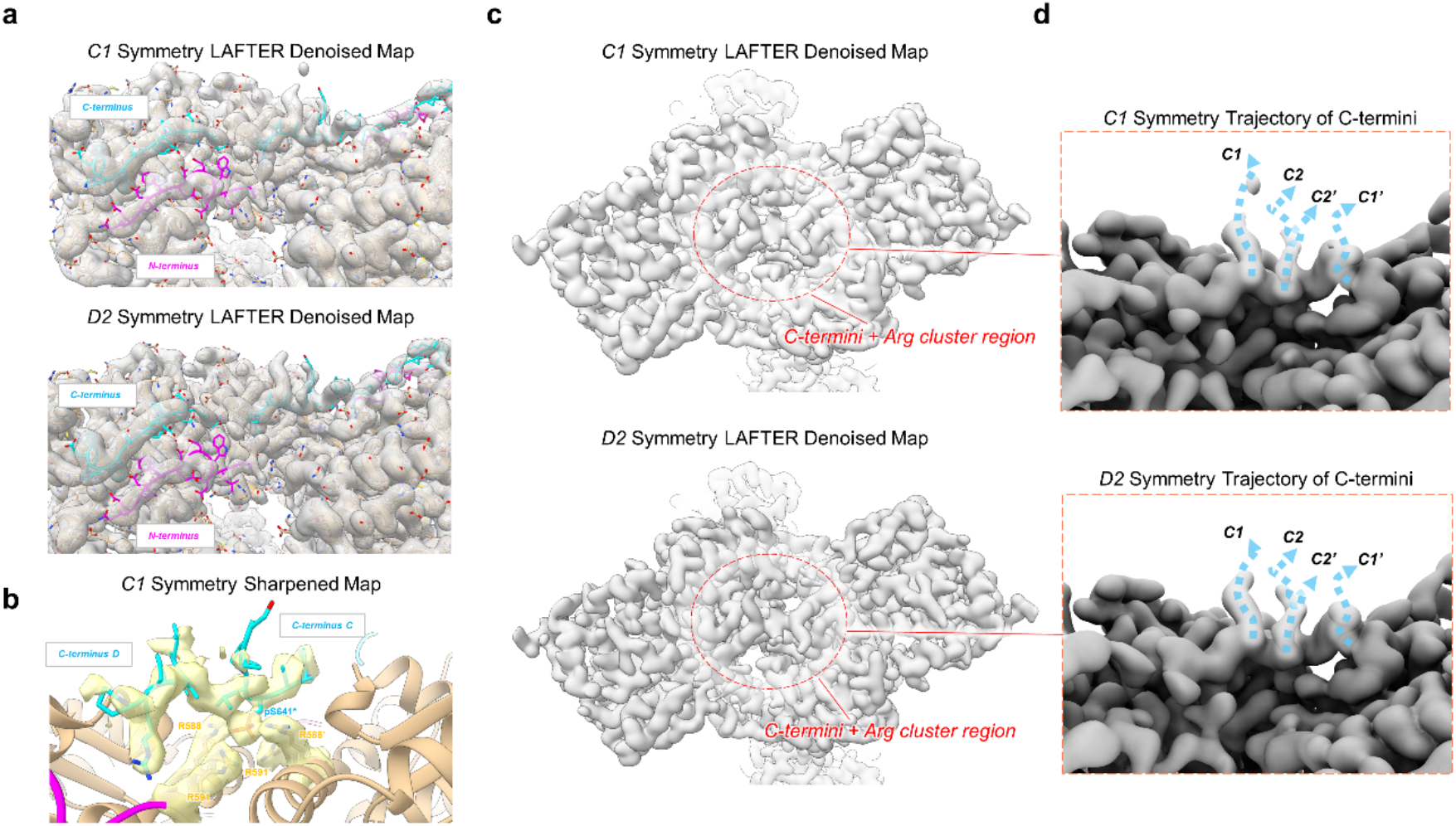
Modelling of the N- and C- termini of the inhibited/T state of the GYS1:GYG1 complex. **a**, Fitting of the N- and C- termini model into the *C1* and *D2* symmetry LAFTER denoised maps. **b**, Fitting of the phosphorylated C- termini model into the sharpened *C1* symmetry map. **c**, Views of the regulatory dimeric interface of the C1 and D2 symmetry LAFTER maps. The phosphorylated C-termini region density is symmetric in both maps. **d**, Predicted directions of the phosphorylated C-termini in C1 and D2 symmetry LAFTER denoised maps. The C-termini are predicted to continue away from the dimeric regulatory interface from two adjacent but different locations.

**Extended Data Fig. 5.**
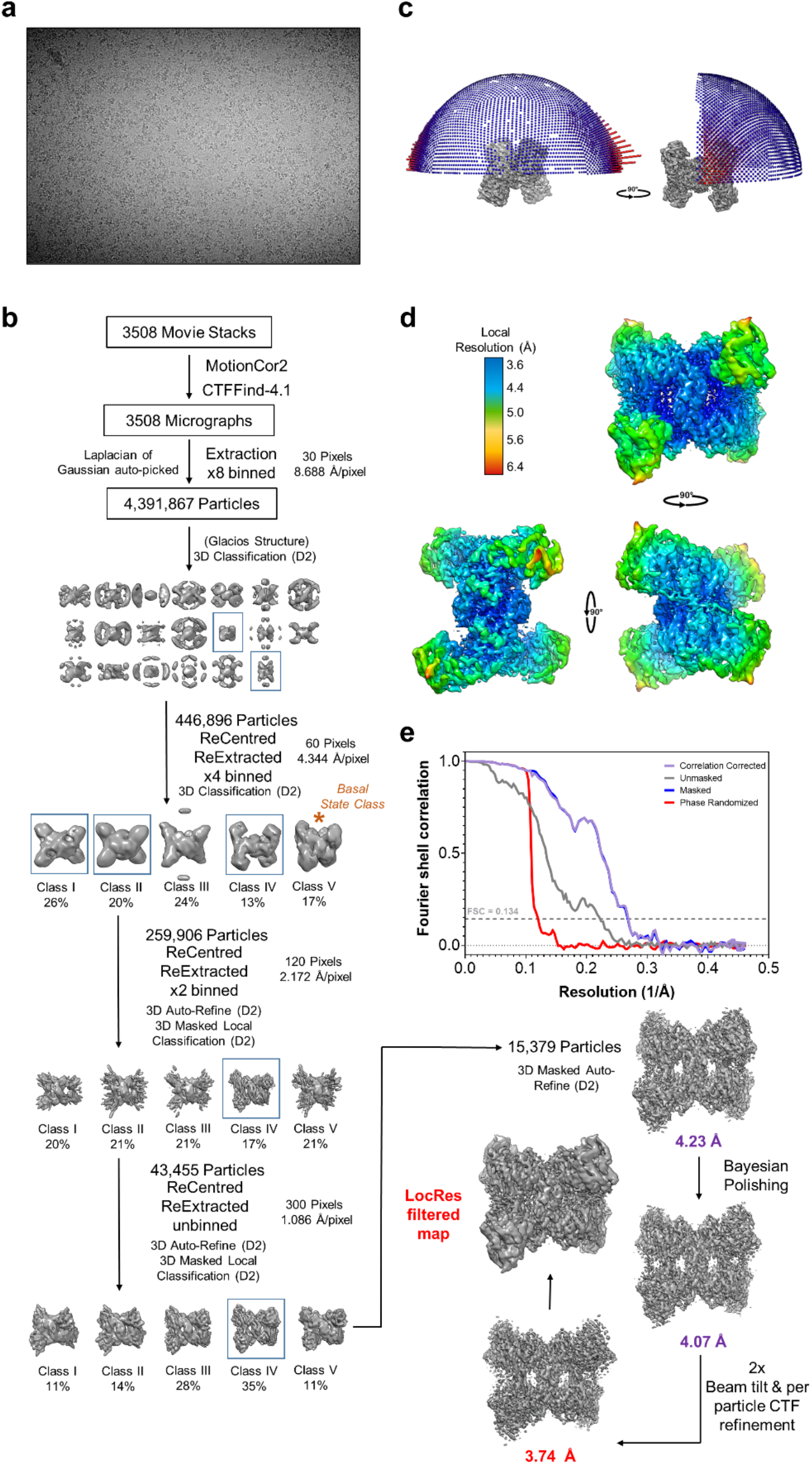
Image processing workflow of the GYS1:GYG1^ΔCD^+Glc6P activated state. **a**, Representative K3 micrograph of the GYS1:GYG1 ^ΔCD^+Glc6P activated state. **b**, Processing flow chart of the GYS1:GYG1 ^ΔCD^+Glc6P activated state. **c**, Angular distribution of the 3.74 Å GYS1:GYG1 ^ΔCD^+Glc6P activated state map. **d**, Local resolution variation of the 3.74 Å GYS1:GYG1 ^ΔCD^+Glc6P activated state map. **e**, FSC curve of the 3.74 Å GYS1:GYG1 ^ΔCD^+Glc6P activated state map.

**Extended Data Fig. 6.**
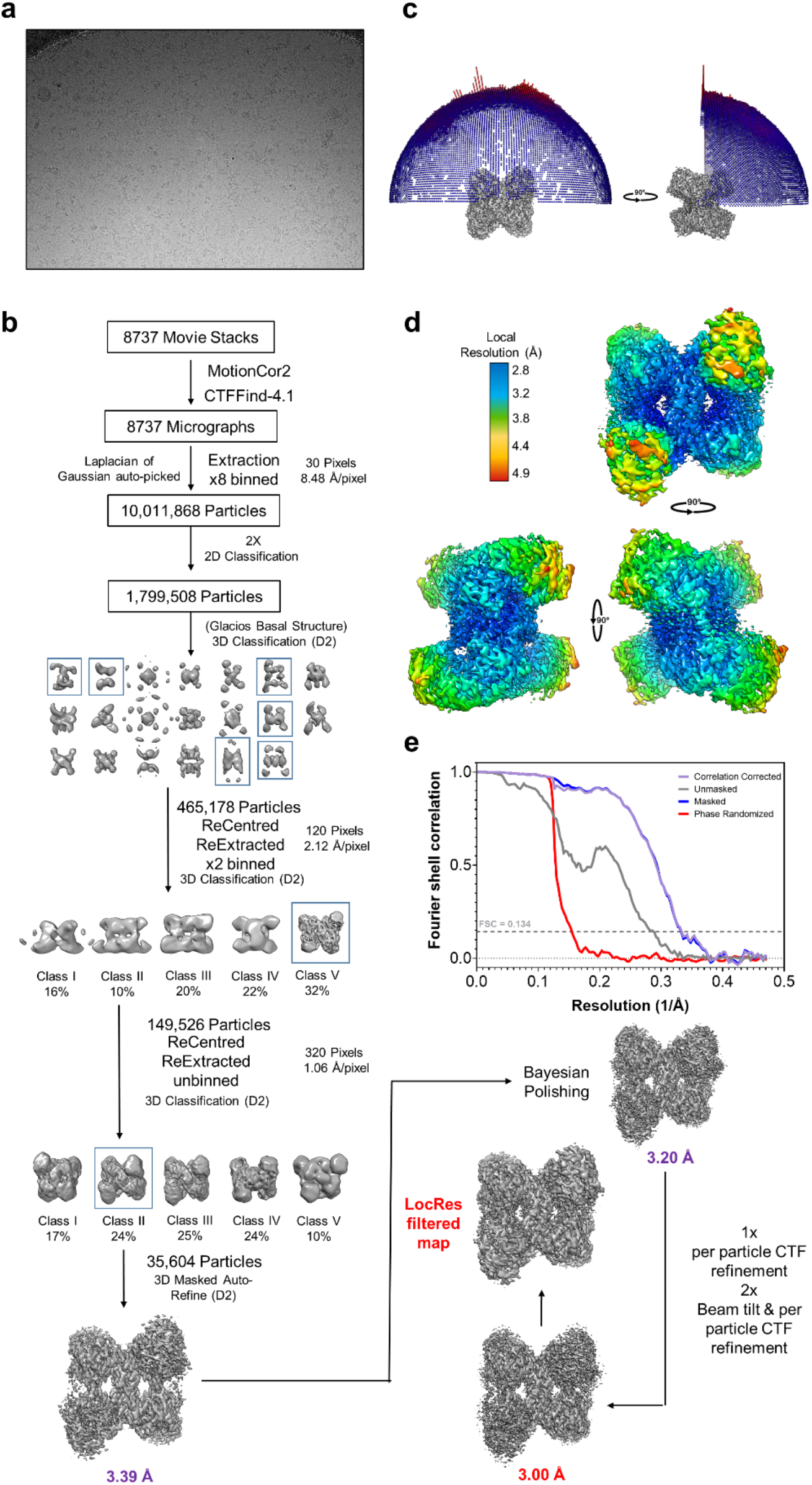
Image processing workflow of the GYS1:GYG1^ΔCD^+Glc6P+UDP-glc activated state. **a**, Representative K3 micrograph of the GYS1:GYG1 ^ΔCD^+Glc6P+UDP-glc activated state. **b**, Processing flow chart of the GYS1:GYG1^ΔCD^+Glc6P+UDP-glc activated state. **c**, Angular distribution of the 3.00 Å GYS1:GYG1^ΔCD^+Glc6P+UDP-glc activated state map. **d**, Local resolution variation of the 3.00 Å GYS1:GYG1 ^ΔCD^+Glc6P+UDP-glc activated state map. **e**, FSC curve of the 3.00 Å GYS1:GYG1^ΔCD^+Glc6P+UDP-glc activated state map.

**Extended Data Fig. 7.**
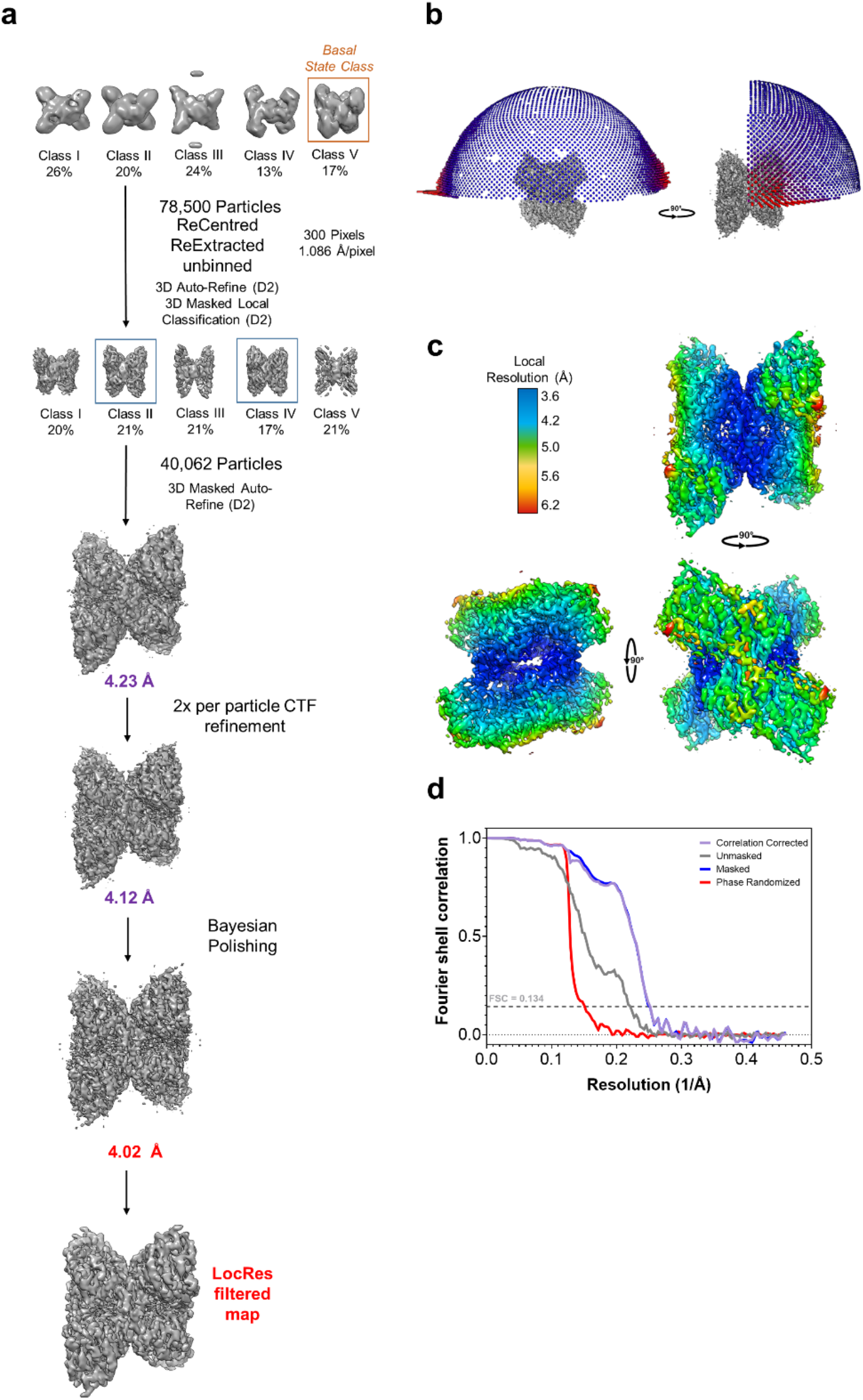
Image processing workflow of the GYS1:GYG1^ΔCD^+Glc6P inhibited-like state. **a**, Representative K3 micrograph of the GYS1:GYG1 ^ΔCD^+Glc6P inhibited-like state. **b**, Processing flow chart of the GYS1:GYG1 ^ΔCD^+Glc6P inhibited-like state. **c**, Angular distribution of the 4.02 Å GYS1:GYG1 ^ΔCD^+Glc6P inhibited-like state map. **d**, Local resolution variation of the 4.02 Å GYS1:GYG1^ΔCD^+Glc6P inhibited-like state map. **e**, FSC curve of the 4.02 Å GYS1:GYG1 ^ΔCD^+Glc6P inhibited-like state map.

**Extended Data Fig. 8.**
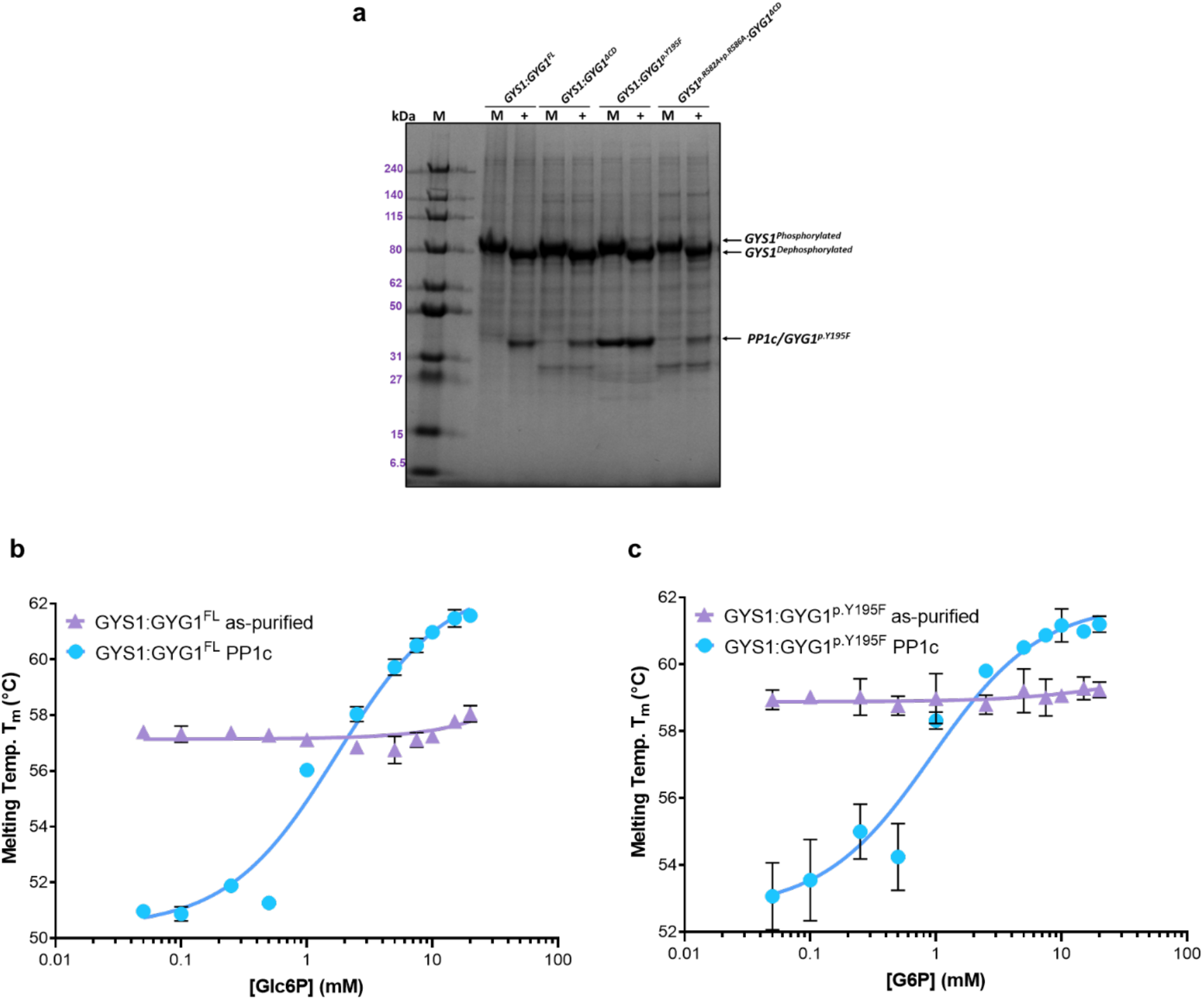
Thermal shift assay of phosphorylated (as purified) versus dephosphorylated (PP1 treated) GYS1:GYG1 complexes in the presence of increasing concentrations of Glc6P. **a**, Gel shift of GYS1:GYG complexes mock (M) or treated with PP1c (+) for 2 hours at room temperature. 5 μg of each complex was loaded and ran on SDS-PAGE. A decrease in the molecular weight of GYS1 after PP1 treatment is apparent. **b**, Thermal shift assay of GYS1:GYG1^FL^ against Glc6P. **c**, Thermal shift assay of GYS1:GYG1^p.Y195F^ against Glc6P. Median melting temperatures and standard deviations are shown (*n* = 4).

**Extended Data Fig. 9.**
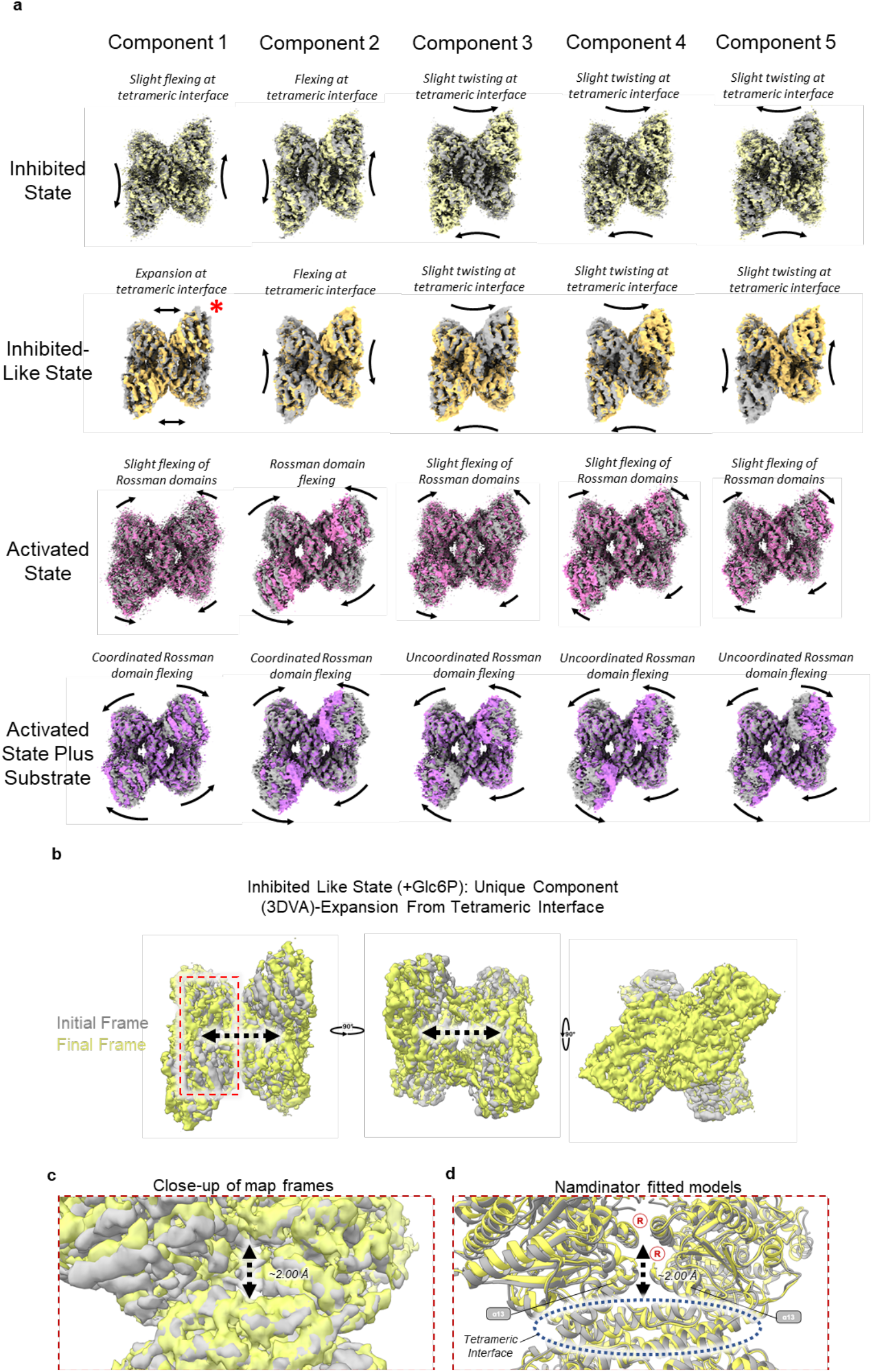
3D variability analysis of the four different states of GYS1 and the unique component of the inhibited like Glc6P bound state. **a**, 3D variability analysis components of all four states of GYS1 reported in this study. Initial and final frames are shown. The unique component of the inhibited like-state is highlighted by a red asterisk. Most movements are either slight flexing at the tetrameric interface or flexing of the N-terminal Rossman domains. **b**, Alignment of initial and final frames showing a global expansion from the central helical tetrameric interface. **c**, Close-up of the frames around the allosteric/G6P binding density **d**, Namdinator fitted models into the initial and final frames showing a clear movement of the alpha-helices 13 from both subunits towards the regulatory helices.

