## SupplementaryFigures for "Molecular basis for the regulation of human glycogen synthase by phosphorylation and glucose-6-phosphate"

[1]Centre for Medicines Discovery, Nuffield Department of Clinical Medicine, University of Oxford, Oxford OX3 7DQ, UK
[2]Rare Disease Research Unit, Worldwide Research and Development, Pfizer Inc., 610 Main Street, Cambridge, MA 02139, USA
[3]Pfizer Inc. Medicine Design, Cambridge, Massachusetts 02139, USA
[4]Discovery Sciences, Worldwide Research and Development, Pfizer Inc., Eastern Point Road, Groton, CT 06340, USA
[5]Present address: Division of Metabolism and Children's Research Center, University Children's Hospital Zürich, University of Zürich, Switzerland
[6]Present address: Biosciences Institute, The Medical School, Newcastle University, Newcastle upon Tyne, NE2 4HH, UK

### Supplementary Figures

**Supplementary Fig. 1:** Sequence alignment of GYS proteins

**Supplementary Fig. 2:** Sequence alignment of the N-terminus and CBM21 domain of each PP1 regulatory subunit and the starch binding domain of *R. oryzae* glucoamylase

**Supplementary Fig. 3:** PTG pulldowns and BN-PAGE replicates

**Supplementary Fig. 4:** Representative thermal shift unfolding curves of each GYS1:GYG1 complex and PTG(CBM21) mutant

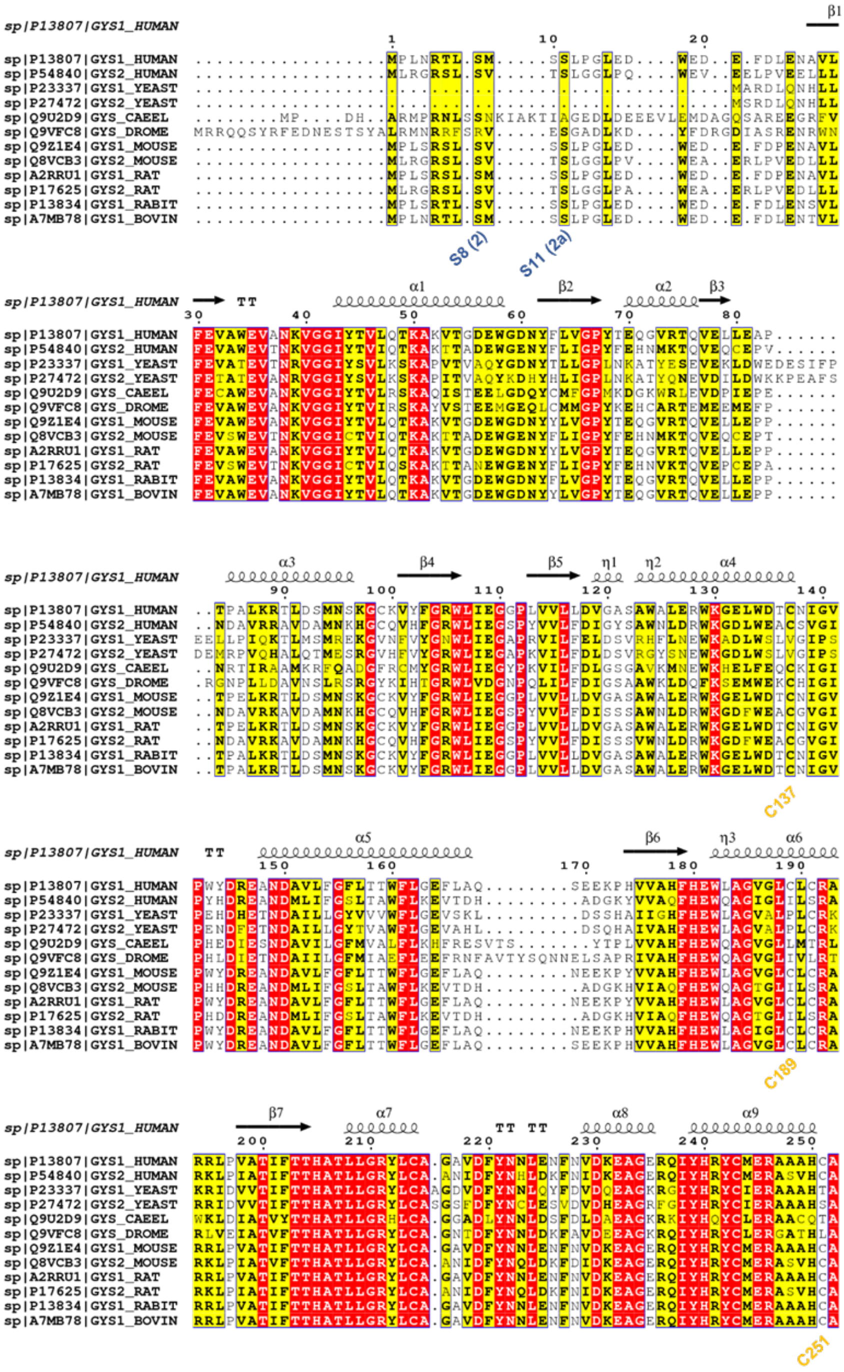
sp|P13807|GYS1_HUMAN
sp|P54840|GYS2_HUMAN
sp|P23337|GYS1_YEAST
sp|P27472|GYS2_YEAST
sp|Q9U2D9|GYS_CAEEL
sp|Q9VFC8|GYS_DROME
sp|Q9Z1E4|GYS1_MOUSE
sp|Q8VCB3|GYS2_MOUSE
sp|A2RRU1|GYS1_RAT
sp|P17625|GYS2_RAT
sp|P13834|GYS1_RABIT
sp|A7MB78|GYS1_BOVIN
S8 (2)
S11 (2a)
C137
C189
C251

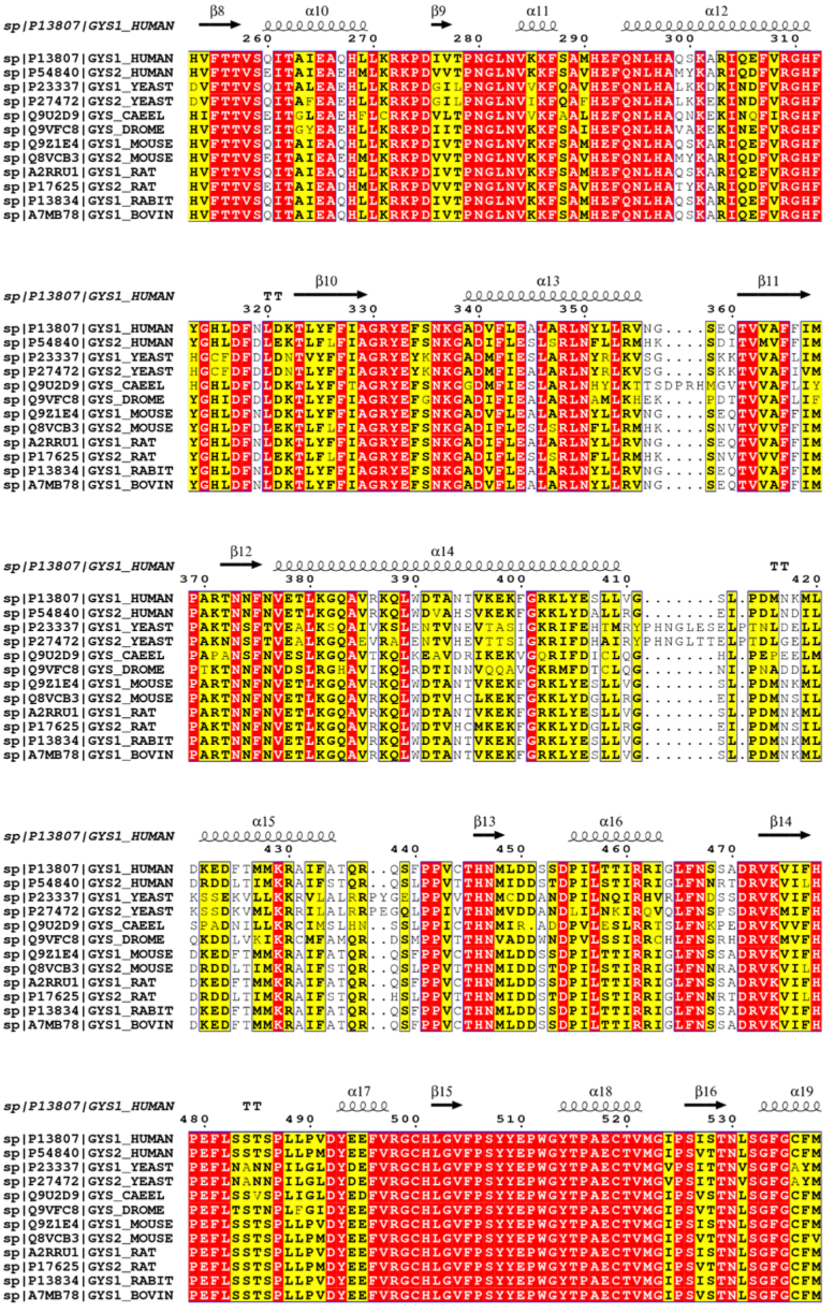
sp|P13807|GYS1_HUMAN
sp|P54840|GYS2_HUMAN
sp|P23337|GYS1_YEAST
sp|P27472|GYS2_YEAST
sp|Q9U2D9|GYS_CAEEL
sp|Q9VFC8|GYS_DROME
sp|Q9Z1E4|GYS1_MOUSE
sp|Q8VCB3|GYS2_MOUSE
sp|A2RRU1|GYS1_RAT
sp|P17625|GYS2_RAT
sp|P13834|GYS1_RABIT
sp|A7MB78|GYS1_BOVIN

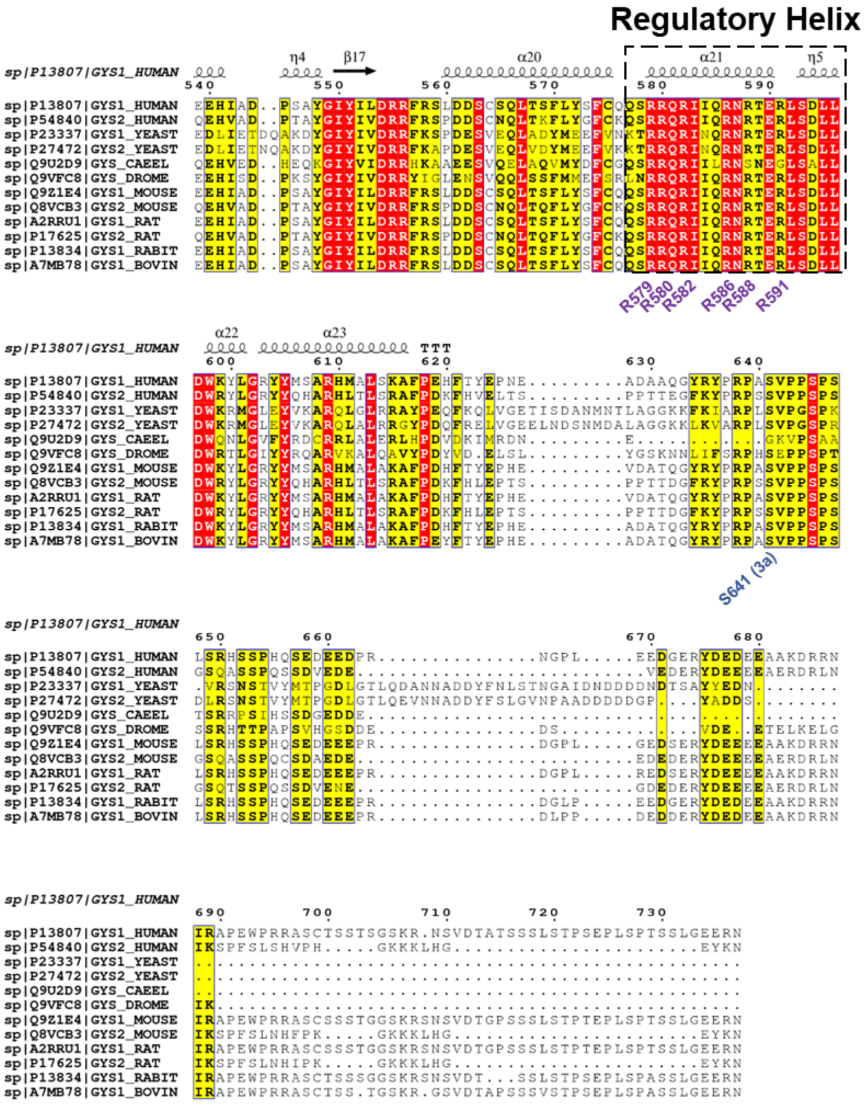


**Supplementary Fig. 1 | Sequence alignment of GYS proteins.** Key regions and residues are labelled.

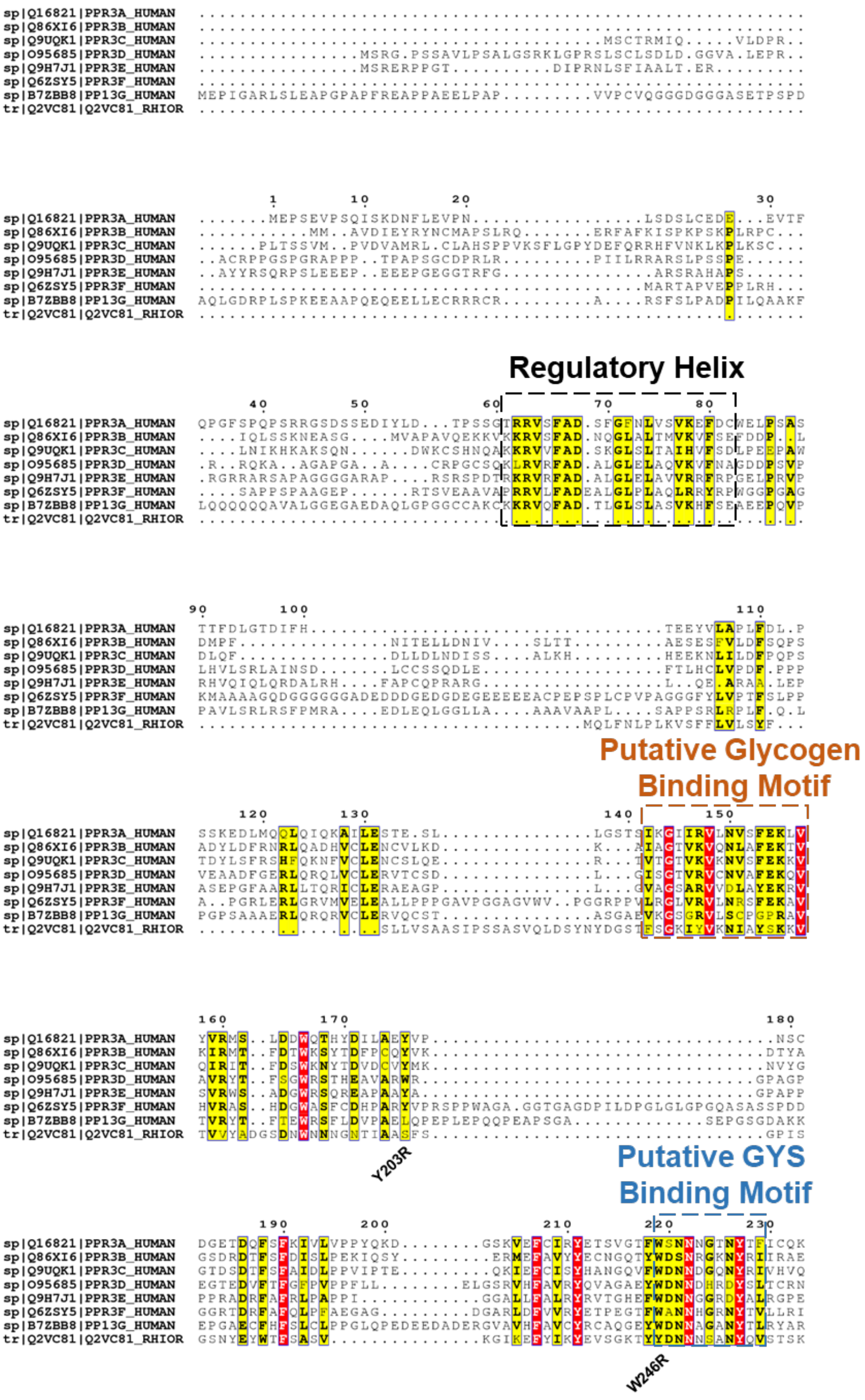


**Supplementary Fig. 2 | Sequence alignment of the CBM21 domain of PP1 regulatory subunit PPP1R3 family and *R. oryzae* glucoamylase.** Only the N-terminus and CBM21 domain of each PP1 regulatory subunit and *R. oryzae* glucoamylase are shown. Key motifs are highlighted.

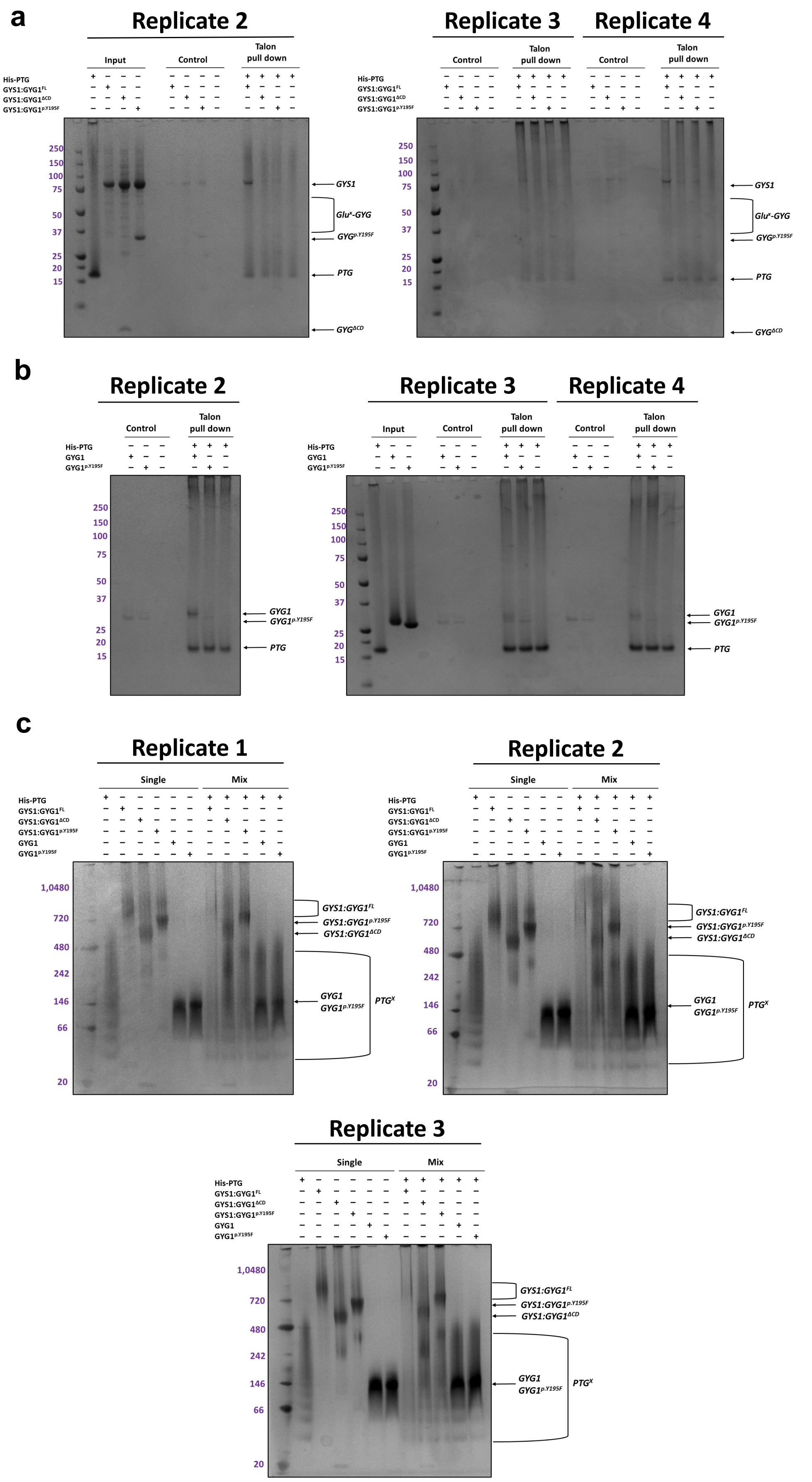

a
Replicate 2
Input
Control
Talon pull down
His-PTG
GYS1:GYG1^FL
GYS1:GYG1^ΔCD
GYS1:GYG1^p.Y195F
250
150
100
75
50
37
25
20
15
GYS1
Glu^x-GYG
GYG^p.Y195F
PTG
GYG^ΔCD
Replicate 3
Replicate 4
b
Replicate 2
Control
Talon pull down
His-PTG
GYG1
GYG1^p.Y195F
GYG1
GYG1^p.Y195F
PTG
Replicate 3
Replicate 4
Input
c
Replicate 1
Single
Mix
His-PTG
GYS1:GYG1^FL
GYS1:GYG1^ΔCD
GYS1:GYG1^p.Y195F
GYG1
GYG1^p.Y195F
1,0480
720
480
242
146
66
20
GYS1:GYG1^FL
GYS1:GYG1^p.Y195F
GYS1:GYG1^ΔCD
GYG1
GYG1^p.Y195F
PTG^X
Replicate 2
Replicate 3

**Supplementary Fig. 3 | PTG pulldowns and BN-PAGE replicates. a**, Coomassie stained SDS-PAGE replicates of PTG(CBM21) pulldowns against GYS1:GYG1$^{FL}$, GYS1:GYG1$^{p.Y195F}$, or GYS1:GYG1$^{\Delta CD}$. **b**, Coomassie stained SDS-PAGE replicates of PTG(CBM21) pulldowns against GYG1$^{FL}$ or GYG1$^{p.Y195F}$. **c**, Blue native PAGE shift replicates of PTG(CBM21) incubated with GYS1:GYG1$^{FL}$, GYS1:GYG1$^{p.Y195F}$, GYS1:GYG1$^{\Delta CD}$, GYG1 alone, or GYG1$^{p.Y195F}$ alone. Complex formation between PTG(CBM21) and GYS1:GYG1$^{FL}$ was inferred from the disappearance of bands compared to other reaction lanes. CBM21 appears as oligomers but behaves as a monomer with some dimer in solution (data not shown).

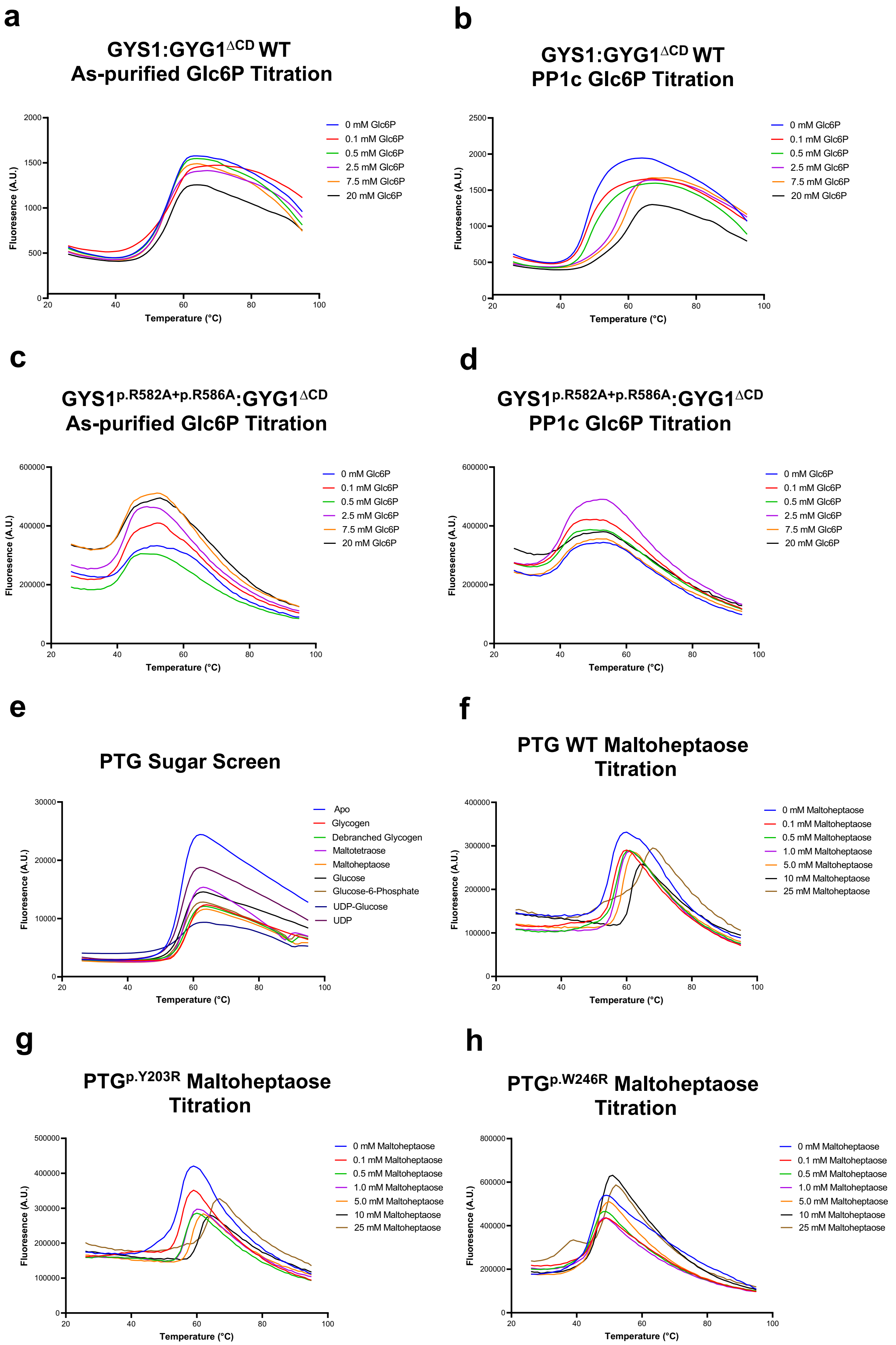
a
GYS1:GYG1ΔCD WT
As-purified Glc6P Titration
b
GYS1:GYG1ΔCD WT
PP1c Glc6P Titration
0 mM Glc6P
0.1 mM Glc6P
0.5 mM Glc6P
2.5 mM Glc6P
7.5 mM Glc6P
20 mM Glc6P
c
GYS1p.R582A+p.R586A:GYG1ΔCD
As-purified Glc6P Titration
d
GYS1p.R582A+p.R586A:GYG1ΔCD
PP1c Glc6P Titration
e
PTG Sugar Screen
Apo
Glycogen
Debranched Glycogen
Maltotetraose
Maltoheptaose
Glucose
Glucose-6-Phosphate
UDP-Glucose
UDP
f
PTG WT Maltoheptaose Titration
g
PTGp.Y203R Maltoheptaose Titration
h
PTGp.W246R Maltoheptaose Titration
0 mM Maltoheptaose
0.1 mM Maltoheptaose
0.5 mM Maltoheptaose
1.0 mM Maltoheptaose
5.0 mM Maltoheptaose
10 mM Maltoheptaose
25 mM Maltoheptaose
Fluoresence (A.U.)
Temperature (°C)

**Supplementary Fig. 4 | Representative thermal shift unfolding curves of each GYS1:GYG1 complex and PTG(CBM21) mutant. a**, GYS1:GYG1$^{\Delta CD}$ WT as-purified, titrated with increasing Glc6P. **b**, GYS1:GYG1$^{\Delta CD}$ WT + PP1c, titrated with increasing Glc6P. **c**, GYS1$^{p.R582A+p.R586A}$:GYG1$^{\Delta CD}$ as-purified, titrated with increasing Glc6P. **d**, GYS1$^{p.R582A+p.R586A}$:GYG1$^{\Delta CD}$ + PP1c, titrated with increasing Glc6P. **e**, PTG(CBM21) screened against various sugars and ligands at 1 mM each. **f**, PTG(CBM21) WT, titrated with increasing maltoheptaose. **g**, PTG(CBM21)$^{p.Y203R}$, titrated with increasing maltoheptaose. **h**, PTG(CBM21)$^{p.W246R}$, titrated with increasing maltoheptaose.
